# Embodied Spatial Knowledge Acquisition in Immersive Virtual Reality: Comparison to Map Exploration

**DOI:** 10.1101/2020.01.12.903096

**Authors:** Sabine U. König, Ashima Keshava, Viviane Clay, Kirsten Rittershofer, Nicolas Kuske, Peter König

## Abstract

Investigating spatial learning in virtual environments allows studying different sources of information under controlled conditions. We built a virtual environment in the style of a European village and investigated spatial knowledge acquisition by experience in the virtual environment and by the use of an interactive map. We tested knowledge of cardinal directions, building-to-building orientation, and judgment of direction between buildings. We find that judgment of directions was more accurate after virtual reality exploration than after map exploration, and the opposite results were observed for knowledge of cardinal directions and relative orientation between buildings. Further, the alignment effect was confined to the map exploration condition. Taken together, our results suggest that the source of spatial exploration differentially influenced spatial knowledge acquisition.

## Introduction

According to theories of embodied and enacted cognition, spatial navigation unfolds in the interaction of the navigator with his or her real-world surroundings (Engel, Maye, Kurthen, & König, 2013; O’Regan & Noë, 2001). Bodily movement provides information about visual, motor, kinesthetic, and vestibular changes, which are essential for spatial cognition (Grant & Magee, 1998; Riecke et al., 2010; Ruddle, Volkova, & Bülthoff, 2011; Ruddle, Volkova, Mohler, & Bülthoff, 2011; Waller, Loomis, & Haun, 2004). Interaction with the environment is a crucial part of acquiring spatial knowledge and is therefore becoming increasingly considered for spatial navigation research (Gramann, 2013).

When acquiring knowledge of a large-scale real-world environment, people combine direct experience during exploration with spatial information from indirect sources such as cartographical maps (Richardson, Montello, & Hegarty, 1999; Thorndyke & Hayes-Roth, 1982). While moving in an environment, people develop knowledge about landmarks and their connecting routes, which is rooted in an egocentric reference frame (Meilinger, Frankenstein, & Bülthoff, 2013; Montello, 1998; Shelton & McNamara, 2004; Siegel & White, 1975; Thorndyke & Hayes-Roth, 1982). Through the integration of this spatial knowledge, a maplike mental representation called survey knowledge may develop (Montello, 1998; Siegel & White, 1975). The acquisition of survey knowledge is supported by using maps and is thought to be coded in an allocentric reference frame (Meilinger et al., 2013; Montello, Waller, Hegarty, & Richardson, 2004; Taylor, Naylor, & Chechile, 1999; Thorndyke & Hayes-Roth, 1982). Previous studies have also found that spatial knowledge acquired from a cartographical map is learned with respect to a specific orientation, whereas orientation specificity is much less consistent in spatial knowledge gained by direct experience (Burte & Hegarty 2014; Evans & Pezdek 1980; McNamara 2003; Meilinger, Riecke, & Bülthoff, 2006; Montello et al. 2004; Presson & Hazelrigg 1984; Shelton & McNamara 2001; Sholl, Kenny, & DellaPorta, 2006). Thus, the acquired spatial knowledge is shaped by the employed source of spatial information, but which spatial knowledge of a large-scale environment derives from direct experience or indirect sources such as maps is still not fully understood.

Investigations of spatial knowledge acquisition in large-scale real-world settings are challenging to perform because of complexity and problems of reproducibility. The rapid progress of technology allows for the design of immersive virtual realities (VRs) that provide the possibility to reduce the gap between classical lab conditions and real-world conditions (Coutrot et al., 2019; Jungnickel & Gramann, 2016). In VR, modern head-mounted displays (HMDs) provide the user with a feeling of presence and immersion in the virtual environment. These VR environments might be considered as “primary” spaces (Montello et al., 2004; Presson & Hazelrigg, 1984), which are directly experienced environments instead of environments indirectly experienced through, for example, maps. Important for a direct experience of the environment is the amount of naturalistic movement. While exploring the VR environment, different degrees of translational and rotational movements, up to walking and body turns in a large, real indoor space or on an omnidirectional treadmill, are realized. These means push the limits of VR and provide sensory information closer to real-world conditions (Darken, Cockayne, & Carmein, 1997; Kitson, Prpa, & Riecke, 2018; Kitson, Riecke, Hashemian, & Neustaedter, 2015; Liang, Starrett, & Ekstrom, 2018; Nabiyouni, Saktheeswaran, Bowman, & Karanth, 2015; Riecke et al., 2010; R. A. Ruddle & Lessels, 2006; R. A. Ruddle, Payne, & Jones, 1999; R. A. Ruddle, Volkova, & Bülthoff, 2011). However, sensory information gathered by experience in a VR environment is mainly visual and in varying degrees vestibular, whereas navigation in the real world is a multisensory process that includes visual, auditory, motor, kinesthetic, and vestibular information (Montello et al., 2004). Another difference between virtual and real environments is the size of the environment that is used to investigate spatial learning. Virtual environments range from containing a few corridors to complex virtual layouts and virtual cities (e.g. Coutrot et al., 2018; Ehinger et al., 2014; Goeke, König, & Gramann, 2013; Gramann, Müller, Eick, & Schönebeck, 2005; Mallot, Gillner, Van Veen, & Bülthoff, 1998; Starrett, Stokes, Huffman, Ferrer, & Ekstrom, 2019; Zhang, Zherdeva, & Ekstrom, 2014). However, motion sickness and the required time for spatial exploration restrict environment size such that it is still small compared to real-world environments. Nevertheless, experiments in VR offer the possibility to investigate spatial learning under controlled conditions, also considering more embodied interaction than classic lab conditions with immersive setups. In summary, despite the differences with respect to real-world navigated environments, immersive VR environments might be viewed as directly experienced environments that allow comparison of spatial knowledge acquisition by direct and indirect sources under controlled conditions with an increased resemblance to spatial learning in the real-world.

In the present paper, we therefore use a virtual village named Seahaven (Clay, König, & König, 2019) and an interactive map of the same environment (König et al., 2019) for a comparison of spatial knowledge acquisition by experience in an immersive VR or by using cartographic material, respectively. For this purpose, one group of participants explored the virtual environment in the VR and another group of different participants used the interactive map for spatial learning. Seahaven consists of 213 buildings and covers an area of 21.6 hectares. Thus, it is a relatively large and complex virtual environment. With the help of an HTC Vive headset, participants had an immersive experience of the environment from a pedestrian perspective during the free exploration in VR. Turning in the immersive VR was realized by bodily turning on a swivel chair in real world and forward movement with the help of a handheld controller. Both movements resulted in corresponding visual changes in the virtual environment. While participants explored the village, we investigated their viewing behavior by measuring eye - and head movements (Clay et al., 2019). During the free exploration of the interactive map of Seahaven, participants were provided with a two-dimensional north-up city map from a birds-eye perspective. Additionally, participants could view screenshots of the buildings of the virtual village with the help of the interactive feature (König et al., 2019). Therefore, we here investigate spatial learning with different sources of the same virtual environment.

After the free exploration of Seahaven, we tested a set of three spatial tasks to investigate, which spatial knowledge is acquired via the different sources. The choice of tasks in the present study was motivated by previous studies investigating the influence of the feelSpace augmentation device, which gives information of cardinal north via vibrotactile information (Kärcher, Fenzlaff, Hartmann, Nagel, & König, 2012; Kaspar, König, Schwandt, & König, 2014; König, Schumann, Keyser, Goeke et al., 2016; Nagel, Carl, Kringe, Märtin, & König, 2005). In those studies, participants who trained with the feelSpace device verbally reported various alignment effects in allocentric reference terms. Due to these reports, we designed three spatial tasks to investigate the learning of allocentric knowledge. The absolute-orientation task evaluated knowledge of orientations of single buildings with respect to cardinal directions. The relative orientation task evaluated the relative orientation of two buildings, and the pointing task investigated judgments of straight-line inter-building directions (König, Goeke, Meilinger, & König, 2017). To investigate intuitive and slow deductive cognitive processes (dual-process theories) that might be used for solving the spatial tasks, we tested all tasks in two response-time conditions, a response window that required a response within 3 seconds and an infinite time condition that allowed unlimited time for a response. All participants, equally in both groups, performed three consecutive full measurement sessions. Additionally to the three tasks, we explored whether spatial-orientation strategies based on egocentric or allocentric reference frames that are learned in everyday navigation, measured with the Fragebogen Räumlicher Strategien (FRS, translated as the “German Questionnaire of Spatial Strategies”) (Münzer, Fehringer, & Kühl, 2016b; Münzer & Hölscher, 2011), were related to the learning of spatial properties tested in our tasks after exploring a virtual village.

Therefore, our research question for the present study is whether and how spatial learning in a virtual environment is influenced by different sources for spatial knowledge acquisition, comparing experience in the immersive virtual environment with learning from an interactive city map of the same environment.

## Methods

### Participants

In this study, we measured spatial knowledge acquisition of 26 participants who explored the virtual environment in an immersive VR repeatedly over the course of three sessions. Participants who suffered from severe motion sickness in their first session were not included in repeated sessions and are thus not included in the 26 participants mentioned above. Out of these 26 participants, four had to be excluded because of technical problems during their measurements. This resulted in 22 participants (11 females, mean age of 22.9 years, SD = 6.7) who explored our virtual environment Seahaven with experience in the immersive VR. Thesewere compared to 22 different participants (11 females, mean age of 23.8 years, SD = 3.1) who explored an interactive map of the virtual environment (König et al., 2019). This matches the target of 22 valid subjects per group as determined by Power analysis (G*Power 3.1, effect size 0.25, determined by a previous study, power 0.8, significance level 0.05, contrasts for ANOVA: repeated measures, within-between interaction).

All participants performed three repeated full sessions, each including a 30-minute exploration phase either in immersive VR or with the interactive map; both groups were tested with the same spatial tasks repeatedly after each exploration phase (Table 1). The three sessions were conducted within a period of ten days. The data that we report here are based on an overall exploration time of 90 minutes and the accuracy of the spatial tasks of the final third session for each participant in both groups. At the beginning of the experiment, participants were informed about the purpose and procedure of the investigation and gave written informed consent. Each participant was either reimbursed with nine euros per hour or earned an equivalent amount of ‘‘participant hours’’, which are a requirement in most students’ study programs at the University of Osnabrück. Overall, each session took about two hours. The ethics committee of the University of Osnabrück approved the study in accordance with the ethical standards of the Institutional and National Research Committees.

**Table 1:** Experiment Procedure

|  | Single Steps in the Experiment | Phases of the Experiment |
| --- | --- | --- |
| 1 | Subject information, written informed consent | Introductory phase (45 minutes) |
| 2 | Response training |  |
| 3 | Spatial tasks instructions |  |
| 4 | Task training with example trials of all tasks in both time conditions |  |
| 5 | Introduction of VR village and how to move in VR or introduction of the interactive city map |  |
| 6 | Setup of VR headset and eye-tracker calibration for VR group |  |
| 7 | Movement practice on small VR practice island for VR group |  |
| 8 | Free exploration directly moving in Seahaven including recalibration of the eye tracker or free exploration of the interactive city map of Seahaven | Exploration phase (30 minutes) |
| 9 | Spatial tasks (absolute-orientation, relative-orientation, and pointing tasks) in two time conditions (3 seconds or infinite time condition) | Test phase (45 minutes)<br>(same for VR and Map group) |
| 10 | FRS questionnaire | Questionnaire phase (5 minutes) |

### VR Village Design

To minimize the gap between the investigation of spatial tasks under laboratory conditions and in real-world environments, we built a virtual environment (Figure 1) named Seahaven. Seahaven is located on an island to keep participants in the village area. Seahaven was built in the Unity game engine using buildings, streets, and landscape objects acquired from the Unity asset store. Seahaven consists of 213 unique buildings, which had to be distinguishable for the spatial tasks and thus have different styles and appearances. The number of buildings sharing a specific orientation toward the north was approximately equally distributed in steps of 30° (0° to 330°). The street layout includes smaller and bigger streets and paths resembling a typical European small village without an ordered grid structure and specific districts. In contrast to a rectangular grid design, this avoids strategies like counting crossings and turns and matches the style of the environment our participants lived in at the time of the study. Seahaven would cover about 21.6 hectares in real-world measure and is designed for pedestrian use. For a virtual environment, Seahaven is relatively large and complex compared to many VR environments used for spatial-navigation research (e.g. Goeke et al., 2013; Gramann et al., 2005; Riecke, Veen, & Bülthoff, 2002; Ruddle, Volkova, & Bülthoff, 2011). Its effective size was limited through the time needed to explore it and because more time spent in the VR increases the risk of motion sickness. As the sun’s position is a primary means to infer cardinal directions in natural surroundings, we implemented this cue given by the sun’s position using a light source in the virtual environment. During the exploration in VR, information about cardinal directions could be deduced from the trajectory of a light source representing the sun over the course of one virtual day. The sun moved on a regular trajectory from East-to-West with an inclination matching the latitude of Osnabrück. The light source’s position indicated a sunrise in the east at the beginning of the exploration, the sun’s trajectory during the day, and the sunset in the west at the end of the same exploration session. As the sun’s position in the sky changed during the session, the shadows of the buildings and other objects in the virtual village changed in relation to the sun’s position as in a real environment.

**Figure 1:**
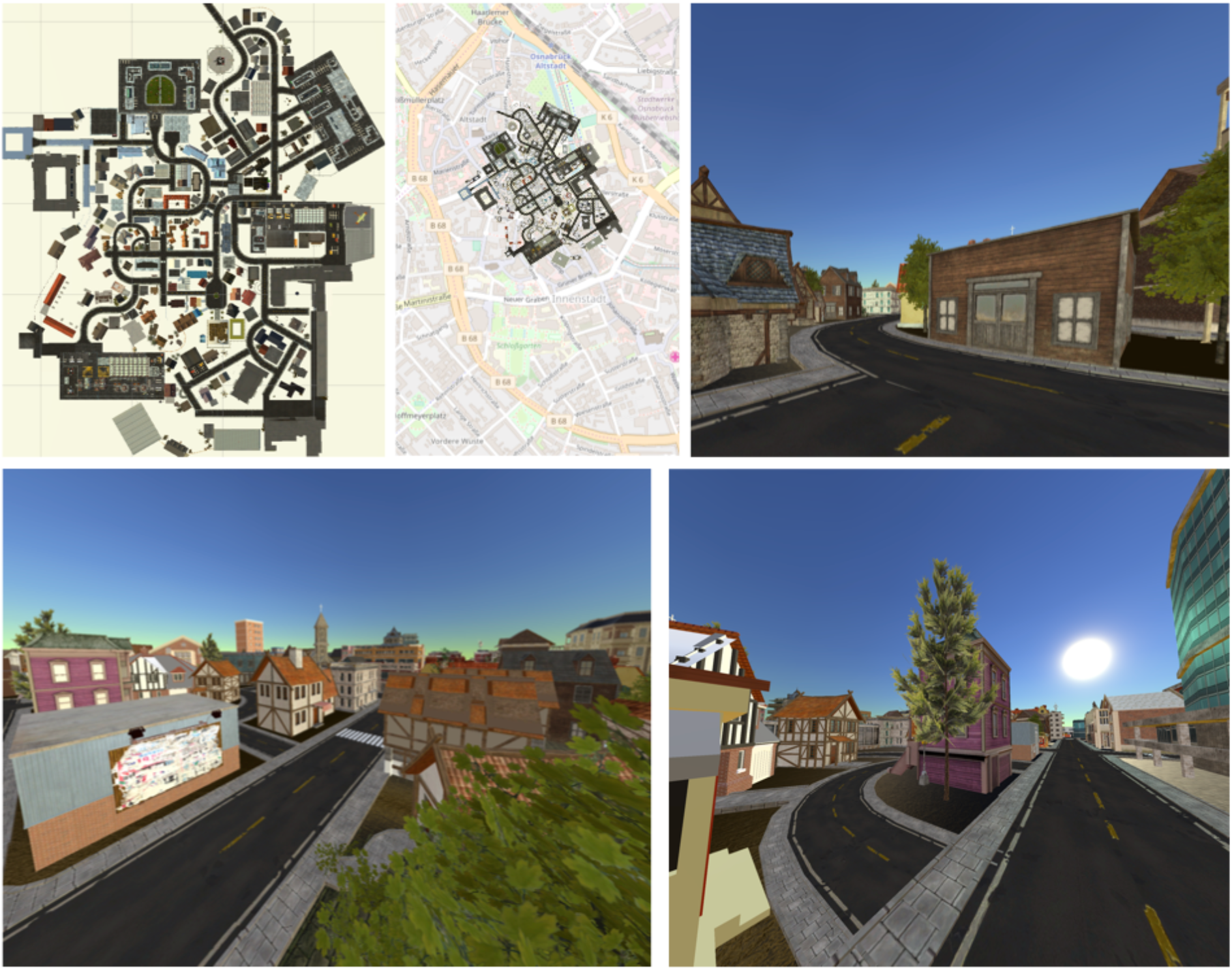
Top-left: Map of the virtual environment. Top-middle: Village map of the virtual city Seahaven overlaid onto the city map of central Osnabrück. Top-right and lower row: The remaining photographs show three examples of views in the village from a pedestrian or oblique perspective.

### Interactive Map Design

The interactive map of the virtual environment resembled a traditional city map with a north-up orientation and a bird’s-eye view (Figure 1, top left). It was implemented using HTML, jQuery, and CSS. By adding an interactive component, participants were also provided with the screenshots of front-on views of 193 buildings in Seahaven from a pedestrian perspective that were used as spatial task stimuli (see below). To view the buildings’ screenshots, participants moved over the map using a mouse. When hovering over a building, a red dot appeared on one side of the respective building. The red dot indicated the side of the building that was displayed in the screenshot. By clicking on the building, the respective screenshot of the building’s façade was displayed.

### Stimuli

The stimuli for the spatial tasks were front-on screenshots of 193 buildings of the overall 213 buildings in Seahaven (examples are shown in Figure 2) and the same for both exploration groups. The screenshots were taken at random times during the day in the virtual village, specifically avoiding consistent lighting conditions that could have been used for orientation information. In the spatial tasks, we compared the orientations of buildings towards north (absolute orientation task) or the relative orientation of two buildings (relative orientation task). As the orientation of buildings can be ambiguous, we took the facing direction of the buildings, which is the direction from which the photographer took the screenshots, as our defined orientation of the buildings (Figure 2). The photographer took the screenshots in the virtual environment from a pedestrian viewpoint that would reflect an approximately 5-meter distance to the corresponding building from a position on a street or path in the VR village. For some buildings, this was not possible, so they were excluded as stimuli. Furthermore, a few buildings were overly similar in appearance and therefore had to be excluded as well. All screenshots were shown in full screen with a resolution of 1920 × 1080 pixels on one screen of a six-screen monitor. For the prime stimuli in the relative orientation and pointing tasks, we used the screenshots of 18 buildings that were most often viewed in a VR pilot study. These prime buildings’ orientations were equally distributed over the required orientations from 0° to 330° to cardinal north, increasing in steps of 30°, and were distributed evenly across the village. Each prime building was used twice in the tasks. The screenshots that were used as target stimuli were only used once.

**Figure 2:**
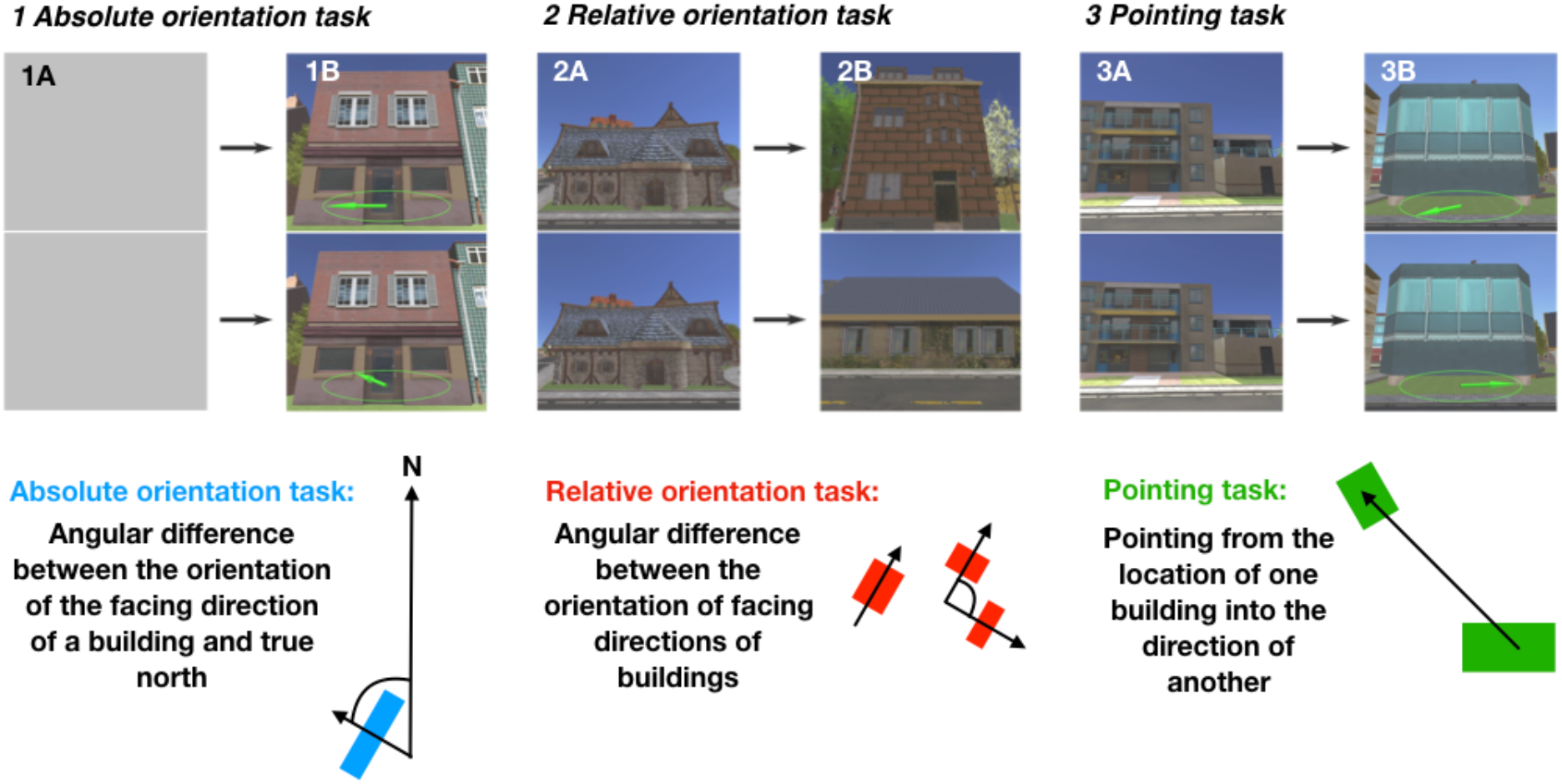
Design of the spatial tasks, depicting example trials of the spatial tasks on the top line and schemata of the tasks in the bottom line: absolute orientation task in blue (1), relative orientation task in red (2), and pointing task in green (3). First, a prime stimulus (A) is shown for 5 seconds in the relative and pointing tasks, which is substituted by a gray screen in the absolute orientation task to fit the trial layout in the other tasks. Then, a target stimulus (B) is shown until button press either in a maximum of 3 seconds or unlimited time to respond. In the absolute orientation task, participants were required to choose the compass needle depicted on the stimuli that correctly pointed to the north. In the relative orientation task, they were required to select the target building that had the same orientation as the prime building. In the pointing task, participants were required to choose the target stimulus in which the compass needle pointed correctly to the location of the prime building (adapted from König et al., 2019) In the schemata, the arrows through the blue and red squares (buildings) depict the facing directions of the respective buildings.

### Spatial Tasks

In our study, we wanted to investigate the learning of allocentric knowledge in terms of cardinal directions, relative orientation of two objects, and straight-line inter-object directions. Therefore, we designed three spatial tasks. Our choice of stimuli for our spatial tasks was motivated by a previous study, which examined the same tasks (König et al., 2017). There, we used photographs of buildings and streets of a real-world city as stimuli as buildings and streets are important landmarks for spatial navigation in real-world cities. For reasons of comparability, we also focused on buildings in the presented study and performed the same spatial navigation tasks with screenshots of buildings of the VR.

Participants were required to perform the same three spatial tasks after VR exploration or map exploration, respectively. All tasks were designed as two-alternative forced choice (2AFC) tasks, requiring the choice of the correct solution out of two possibilities. Thus, the answers were either correct or wrong (Figure 2). In the absolute orientation and the pointing task, the choice options were depicted as compasses of which one compass needle pointed into the correct direction and the other into a wrong direction. These correct and wrong directions deviated from each other in different angular degrees in steps of 30° from 0° to 330°. In the relative orientation task, the two choices were buildings with different orientations, one had the same and, thus, correct orientation as the prime building, the other had a different orientation. Again, angular differences between correct and wrong orientation varied in steps of 30° from 0° to 330°. These angular differences between choice options provided the information from which we calculated the angular differences and the alignment to north.

In the absolute orientation task, participants were required to judge the orientation of single buildings in comparison to the northerly direction (Figure 2, left bottom line). In one trial of the absolute orientation task, two screenshots of the same building were overlaid with a compass; one compass needle pointed correctly toward cardinal north and the other one pointed randomly into another direction deviating from north in steps of 30° (0° to 330°; Figure 2, left top line). Out of the two images of the building, participants were required to choose the image on which the compass needle pointed correctly to cardinal north.

In the relative orientation task, participants judged the orientation of buildings relative to the orientation of another building (Figure 2, middle bottom line). In one trial of the relative orientation task, participants saw a prime building followed by two different target buildings that differed in their orientation in steps of 30° (0° to 330°) from each other (Figure 2, middle top line). Out of the two buildings, participants were required to choose the target building that had the same orientation as the prime building: in other words, the target building whose orientation was closely aligned with the prime building’s orientation.

In the pointing task, participants had to judge the direction from the location of one building to the location of another building (Figure 2, right bottom line). In one trial of the pointing task, a screenshot of a prime building was presented first, followed by two screenshots of the same target building. This target building was depicted twice: once overlaid with a compass needle correctly pointing into the direction of the prime building and once overlaid with a compass needle randomly pointing in another direction deviating from the correct direction in steps of 30° (0° to 330°; Figure 2, right top line). Of these two images of the target building, participants were required to choose the target building with the compass needle that correctly pointed in the direction of the location of the prime building.

All three tasks were performed in two response-time conditions, the 3 seconds condition with a 3 seconds response window and the infinite time condition with unlimited time to respond. This resulted in six different conditions that were presented in blocks. Every block consisted of 36 trials. The number of trials, 216 in total, is a balance between the quality of data and load on participants’ alertness and motivation. The order of blocks was randomized across subjects. For more detailed information, see König et al. (2017, 2019).

### Overview of Experimental Procedure

Our experiment consisted of four major phases, the same for both groups, with only small adjustments for the source of spatial exploration, which are described in detail below (Table 1).

The first was the introductory phase, which lasted approximately 45 minutes. Here, participants were informed about the experiment and gave written informed consent. Further, they performed the response training and received instructions for and an explanation of the spatial tasks. The introduction of the spatial tasks included example trials for all task and time conditions (see section “Response Training” and “Spatial Tasks’ Instructions and Task Training” below). Next, participants in the immersive VR group were introduced to VR and instructed how to move in Seahaven. They were especially informed about the risk of motion sickness. After this, an HTC Vive headset with an integrated eye tracker was mounted, giving an immersive VR experience. With this, participants practiced their movement in VR, which was followed by calibration and validation of the eye tracker. Participants in the map group were instead introduced to the interactive map and instructed how to use the interactive feature.

For the second phase, the exploration phase, VR participants were placed in a predefined place in Seahaven from which they started their free exploration of the virtual village for 30 minutes. Every 5 minutes, the exploration was briefly interrupted for validation of the eye tracker. The map group freely explored the interactive map of Seahaven, also for 30 minutes. For this, participants freely moved on the city map with a mouse and by clicking on a building were provided with screenshots of the building’s front view through the interactive feature (see section “Exploration of Seahaven’s Interactive City Map” and Figure 3).

**Figure 3:**
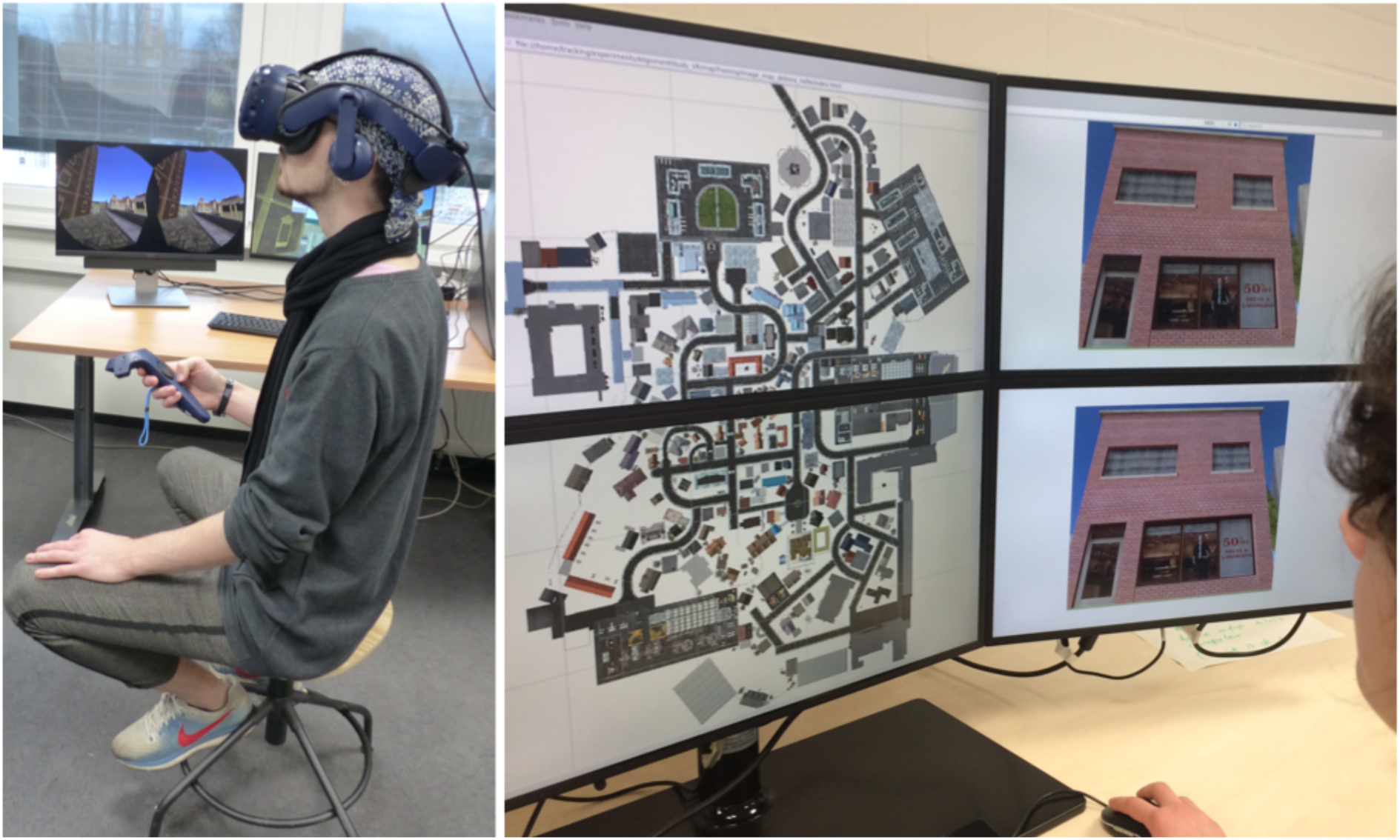
Left: Experimental setup of a participant with the HTC Vive, controller, and swivel chair. Right: Experimental setup of a participant using the interactive map (written informed consent was obtained from the individuals for the publication of these images).

The third (testing) phase lasted for approximately 45 minutes. Here, participants in the VR and map group were tested on the same three spatial tasks (absolute-orientation, relative-orientation, and pointing tasks) in two time conditions (3 seconds and infinite response time condition; see section “Stimuli” and “Spatial Tasks”).

Fourth and finally, all participants filled out a questionnaire on spatial strategies (FRS questionnaire), which concluded the experiment.

### Response Training

To familiarize participants with the 3-second response, the interpretation of the directional compass needle on the screen, and the behavioral responses in the required the 2AFC tasks, each participant performed a response training. Here, the participants were required to compare two compass needles and then select within 3 seconds the compass needle that pointed most straightly upward on the screen. Therefore, one compass needle was presented on one screen and another compass needle on another screen. The screens were placed above each other (Figure 3) and the compass needles pointed in different directions (see Figure 2 with examples of spatial tasks). To choose the compass needle on the upper screen, participants were required to press the “up” button. To select the compass needle on the lower screen, they were required to press the “down” button. On each trial, they received feedback on whether they decided correctly (green frame), incorrectly (red frame), or failed to respond in time (blue frame). The response training was finished when the participants responded correctly without misses in 48 out of 50 trials (accuracy > 95%). This response training ensured that participants were well acquainted with the response mechanism of the 2AFC design that our spatial tasks used.

### Spatial Task Instructions and Task Training

A separate pilot study, in which participants did not know the tasks before the exploration phase, revealed that, during their free exploration, they sometimes focused more on aspects that would not support spatial learning, such as detailed building design. It is known that during spatial learning, paying attention to the environment or the map supports spatial-knowledge acquisition (Montello, 1998). When acquiring knowledge of a new city, knowledge of where important places such as home, work, and a bakery are is crucial for everyday life and thus provides motivation to learn the spatial relations. To support the motivation of spatial exploration in VR, each participant received task instructions and task training before the start of the free exploration time in Seahaven to support intentional spatial learning. Note that none of the subjects of the pilot study is included in the main study and that no subjects of the main study were excluded for reasons of their spatial exploration behavior. The instructions were given using written and verbal explanations accompanied by photographs of buildings in the city of Osnabrück that were used as stimuli in a previous study (König et al., 2017). Participants then performed pre-task training with one example of each spatial task in both time conditions to gain a better insight into the actual task requirements. Except for the stimuli, the pre-tasks exactly resembled the design of the spatial tasks (see sections “Stimuli” and “Spatial Tasks” below). We performed the task training to familiarize participants with the type of spatial knowledge that was later tested in the spatial tasks.

### Exploration of Seahaven in Immersive VR

To display the virtual environment in an immersive fashion, we used an HTC Vive headset (https://www.vive.com/eu/product/; Figure 3, left) with an integrated Pupil Labs eye tracker (https://pupil-labs.com/vr-ar/). Participants were instructed about the virtual village and the potential risk of motion sickness. Additionally, they were informed that they could terminate their participation at any time without giving reasons. During the exploration phase, participants were seated on a swivel chair which allowed for free rotation. For moving in VR, participants were instructed to move forward in the VR environment by touching the forward axis on the HTC Vive’s handheld controller touchpad. To turn, they were required to turn their physical body on the swivel chair in the real world. After these instructions, the HTC Vive headset was mounted, and participants themselves adjusted the interpupillary distance. To support immersion in VR and avoid distractions due to outside noises, participants heard the sound of ocean waves over headphones. Participants started in the VR on a small practice island where they practiced their movements in VR until they felt comfortable. This was followed by the calibration and validation of the eye tracker (see section “Eye Tracking in Seahaven” below). Then, they were placed in the virtual village at the starting position: the same central main crossing for all participants. From there, they freely explored Seahaven for 30 minutes by actively moving within the virtual environment. The participants were informed that it was morning in Seahaven when they began their exploration and that at the end of the 30-minute exploration the sun would be close to setting (see section “VR Village Design”). We validated the usefulness of this approach by asking the participants to turn toward north at the end of the exploration session in VR. Thus, Seahaven provided a virtual environment for immersive free spatial exploration.

### Eye and Position Tracking in Seahaven

During the free exploration in the immersive VR, we recorded the viewing behavior and the position of the participants in the virtual village. For directly measuring eye tracking in VR, we used the Pupil Labs eye tracker, which is plugged into the HTC Vive headset (Clay et al., 2019). To ensure reliable eye-tracking data, we performed calibration and validation of the eye tracker until a validation error below 2° was achieved. We performed a short validation every 5 minutes to check the accuracy, for which slippage due to head movements was possible. If the validation error was above 2°, we repeated the calibration and validation procedure. From the eye-tracking data, we evaluated where participants looked. For this, we calculated a 3D gaze vector and used the ray casting system of Unity, which detects hit points of the gaze with an object collider. With this method, we obtained information about the object that was looked at and its distance to the eye. The virtual environment was differentiated into three categories: buildings, the sky, and everything else. As Unity does not allow the placement of a collider around the light source that we used to simulate the moving sun, we could not investigate hit points with the sun. As buildings were our objects of interest, each building was surrounded by an individual collider. This enabled us to determine how many, how often, for how long, which, and in which order buildings were looked at, giving us information about participants’ exploration behavior. For the position tracking, we recorded all coordinates in Seahaven that a participant visited during the free exploration. Taken together, we measured the walked path and viewing behavior of participants, which enabled us to characterize their exploration behavior in the virtual environment.

### Exploration of Seahaven’s Interactive City Map

For the exploration of the interactive map of Seahaven (Figure 1, top left and Figure 3, right), participants sat approximately 60 cm from a six-screen monitor setup in a 2 × 3 screen arrangement. During the exploration, the two-dimensional map of the virtual environment was presented on the two central screens (Figure 3, right). To explore the map, participants moved over the map using a mouse. When hovering over a building, a red dot appeared on one side of the respective building. By clicking on this building, the interactive component displayed the screenshot of the building twice on the two right screens of the monitor (the same image above each other) (Figure 3, right). How often buildings were clicked on was recorded and later used to determine participants’ familiarity with the stimuli and which buildings were looked at to investigate the part of Seahaven that participants visited. The two screens on the left side of the six-screen monitor setup were not used during exploration or testing. In summary, the interactive map provided participants with a city map of Seahaven and front-on views of the majority of buildings of Seahaven.

### FRS Questionnaire

To compare participants’ abilities in spatial-orientation strategies learned in real-world environments with their accuracy in the spatial tasks after the exploration of a virtual village, participants filled in the FRS questionnaire (Münzer & Hölscher, 2011) at the end of the measurements. The FRS questionnaire imposes self-report measures for spatial orientation strategies learned in real environments. It captures one strategy that is based on an egocentric reference frame and two strategies that are based on an allocentric reference frame (Münzer et al., 2016b; Münzer, Fehringer, & Kühl, 2016a; Münzer & Hölscher, 2011). The “global-egocentric scale” evaluates global orientation abilities and egocentric abilities based on knowledge of routes and directions. The evaluation of allocentric strategies is separated into the “survey scale”, which assesses an allocentric strategy for mental map formation and the “cardinal directions scale”, which evaluates knowledge of cardinal directions. All scales consist of Likert items with a score ranging from 1 (“I disagree strongly.”) to 7 (“I agree strongly.”). Thus, the FRS questionnaire enables us to obtain insights into participants’ preferred use of egocentric or allocentric spatial strategies and to investigate whether spatial abilities learned in the real world are used for spatial learning in a virtual environment.

### Data Analysis Using Logistic Regression for Binary Answers

As we have binary answers in our 2AFC task design, we performed a logistic regression analysis. The modeling was done using R 3.6.3. We used the glm() function from the lme4 package. The 3 models can be formalized as follows using the Wilkinson notation (Wilkinson & Rogers, 1973):

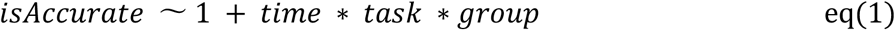

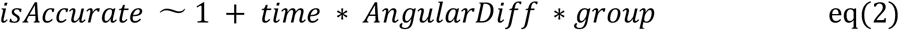

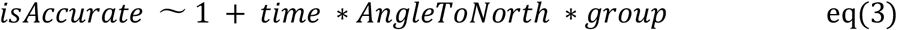

Where “*isAccurate*” is the accuracy, “*time*” codes the 3 seconds or infinite condition, “*task*” codes the absolute, relative, and pointing task, “*group*” codes VR and map exploration group, “*AngularDiff*” codes the angular difference of the two response options, and “*AngleToNorth*” codes the angle of the response to north;

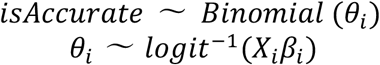

where X is the design matrix and *β* are the estimated parameters of the model.

We modeled the log odds of being accurate for each trial based on task (absolute, relative, or pointing), time (3 seconds or infinite time condition), and exploration group (map or VR) and their interactions. The categorical variables of task, time, and group were coded using simple effect regression coding so that the log odds ratios could be directly interpreted as the main effects from the model coefficients. The model gives log odds ratios for the different levels of each variable with respect to the reference level. Therefore, no separate post-hoc tests are required. We also used the same model for analyzing the angular differences between task choices and alignment to north. Similarly, the log odds of being accurate for the angular differences between task choices and alignment to north were modeled as per eq(2) and eq(3).

Following model fitting, we performed chi-squared tests to compare each of the models to evaluate whether the model parameters perform better than a null model fitted only with an intercept.

Further, we performed an ordinary least squares linear regression for analyzing the distance effect. We used the ols() function of the Python package statsmodels v0.11. The dependent variable was the accuracy of participants for a given prime-target building pair, and the independent variables were the task, time, and group as above.

## Results

In this paper, we investigated the acquisition of spatial knowledge by experience in a large-scale immersive virtual village. In close analogy, a previous study used an interactive map for the spatial exploration of the same virtual environment and reported a preliminary analysis of those data (König et al., 2019). Here, data obtained from both experiments are analyzed in depth using identical procedures (generalized linear models), and fully reported to enable direct comparison between the two sources of spatial learning.

### Results of Exploration Behavior

While participants freely explored the virtual village, we measured their viewing behavior and tracked their positions, which were summed to show their walked path. Analyzing the viewing behavior, we focused on looks on buildings. Here, we were interested in how long a building was viewed in total. Therefore, we calculated the summed dwelling time over the three sessions on a building averaged over subjects (Figure 4, left) and took this as a measure of familiarity of the building in question (see analysis below).

**Figure 4:**
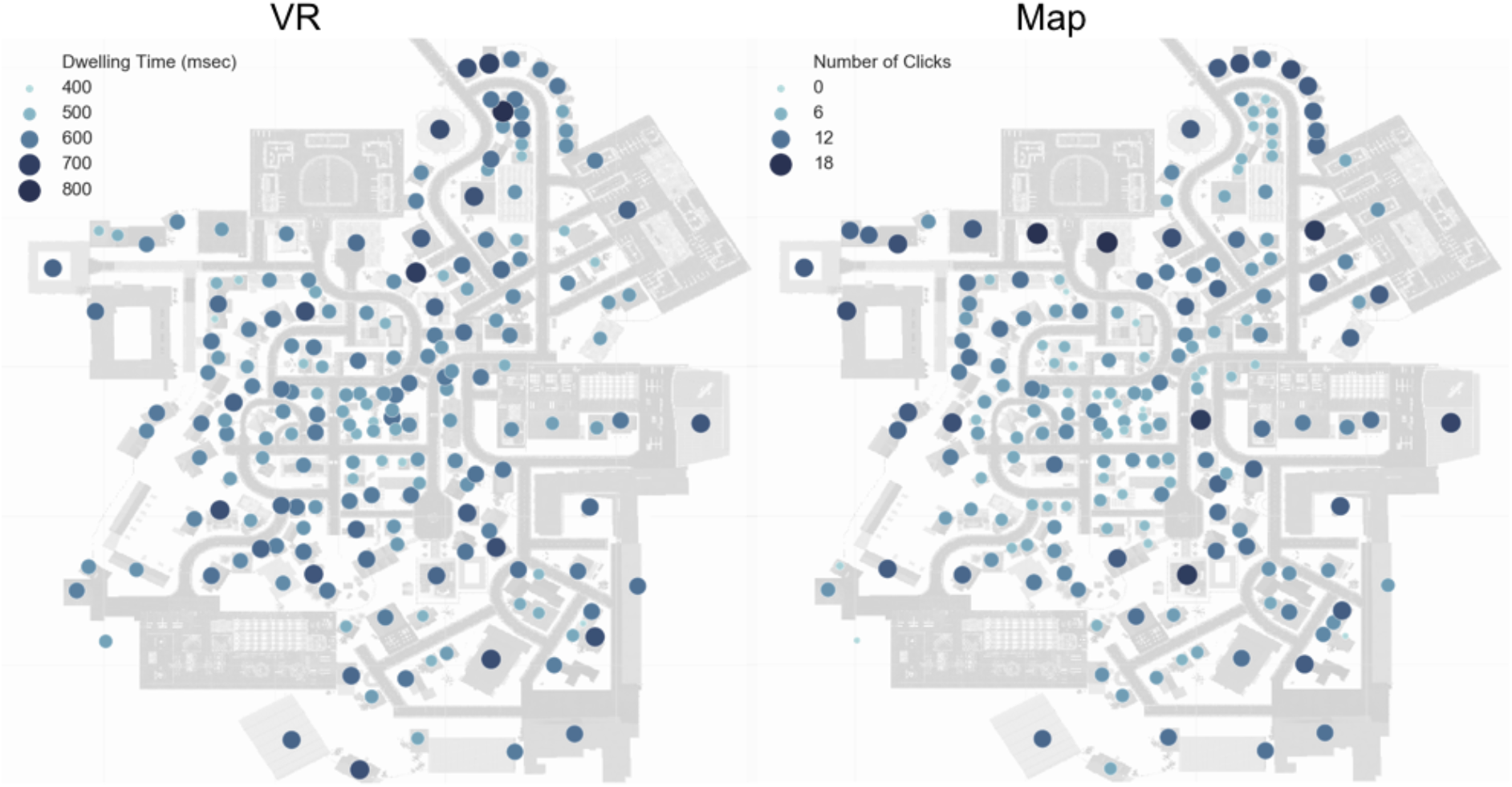
Heat map depicting how long participants looked at a building with exploration in immersive VR (left) and how often participants clicked on buildings with map exploration (right) on a gradient going from deep blue (most) to light blue (least).

For participants, who explored the interactive city map of Seahaven, we measured how often a building was clicked on. Clicking on a building displayed the screenshot of the respective building. We summed the clicks on a building averaged over subjects (Figure 4, right) as a measure of the familiarity of the building after map exploration. Overall, clicks on buildings revealed the parts of Seahaven that were visited during map exploration.

The distribution of the dwelling time on buildings in the village revealed that buildings that were looked at the longest were not centered in a particular village district but were distributed throughout the village. The same held true for the distribution of clicks on buildings after map exploration. For comparison of the distribution of dwelling time on a building after VR exploration to the clicks on buildings when exploring the city map, we performed a Spearman’s rank correlation between dwelling time and the number of clicks on buildings. The result revealed a moderately significant correlation (rho(114) = 0.36, p < 0,001) of buildings, which were explored in the VR and clicked-on buildings during exploration of the city map (Figure 5). The distribution of looked at buildings in VR and clicked on buildings with the interactive map over the virtual village and their correlation suggest that the exploration covered Seahaven well for both exploration sources.

**Figure 5:**
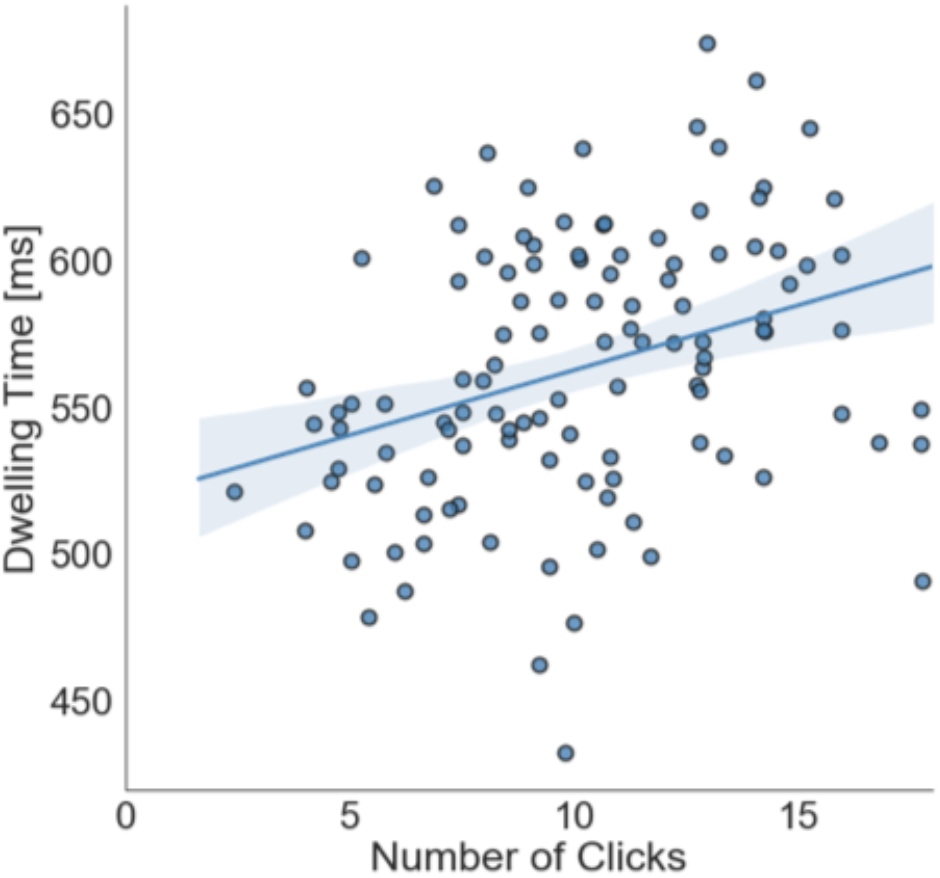
Correlation between dwelling time on buildings with VR exploration (in ms) and number of clicks on buildings with map exploration. Each dot depicts a dwelling time × click combination of a building averaged over subjects. The blue line depicts the regression line and the shaded area the 95% confidence interval.

To investigate the walked path of participants exploring the immersive virtual environment, we analyzed all coordinates in Seahaven which a participant visited and displayed this as a density distribution of the count of the “presence at a location” plotted onto the city map (Figure 6, left). We performed a Pearson correlation between distance from the center of gravity of the city map and density count of presence at a location (log_10_(bin of density count)) and found a weak negative correlation (rho(22340) = −0.12, p < 0.001) (Figure 6, right). The density map of presence at a location investigating the walked path in VR revealed that participants visited all possible locations but that central streets were visited slightly more often than more peripheral paths during participants’ exploration of the virtual environment.

**Figure 6:**
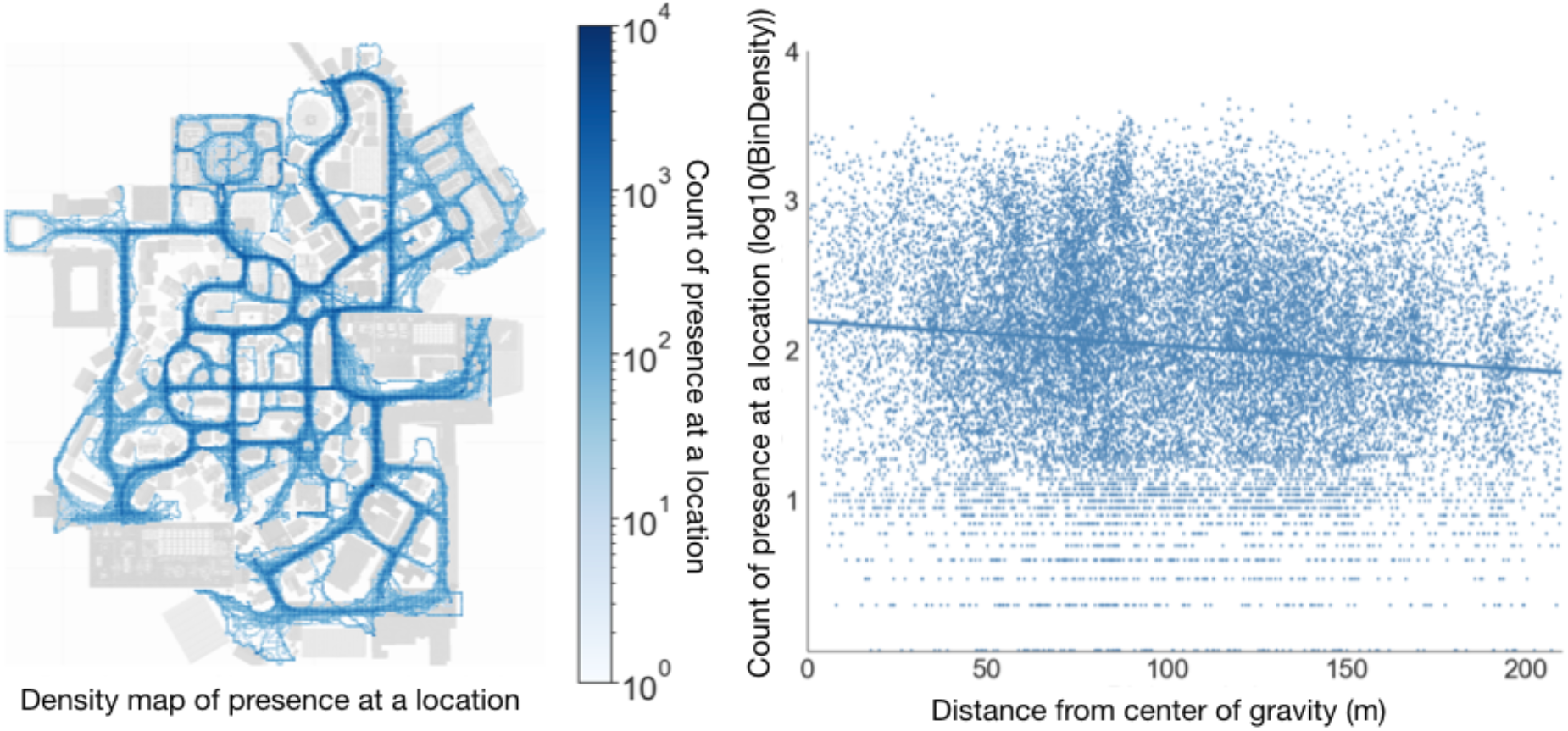
Density distribution of presence at a location after exploration in VR for all participants. Left: depicting count of presence at a location in the virtual environment plotted onto the village map. Right: Correlation between distance from the center of gravity of the city map and density count in bins of presence at a location. Blue dots depict singly visited locations by a participant, and the blue line is the regression line.

### Spatial Task Results

For the performance in the spatial tasks, we evaluated the accuracy of response choices in the 2AFC spatial tasks after experience in the virtual village. We analyzed the accuracy in the different task and time conditions comparing the results after exploration in the immersive VR with the results after exploration of an interactive city map (König et al., 2019). Subsequently, we investigated the influence of how long (dwelling time) a building was looked at (familiarity of buildings), the angular difference between choice options, alignment of stimuli toward north, the distance between tested buildings, and abilities of spatial strategies (FRS questionnaire) on task accuracy.

### Task Accuracy in Different Time Conditions Comparing VR and Map Exploration

Following dual-process theories (Evans, 1984; Evans, 2008; Kahneman & Frederick, 2002), we hypothesized that the accuracy with unlimited time for a response would be higher than with restricted time to respond within 3 seconds. Furthermore, following previous research using this set of tasks (König et al., 2017), we hypothesized that experience in the immersive virtual environment would support action relevant tasks and therefore reveal a higher accuracy for judging straight-line directions between buildings tested in the pointing task as well.

To investigate whether the performance data are significantly different from chance, we performed a binominal test, which revealed a significant difference from chance level (p < 0.01). The mean of task accuracies indeed showed that with unlimited time for response accuracy was in all tasks higher than that with a response within 3 seconds (Table 2). The means also showed that task accuracy was higher in the pointing task (by 3.0%) than absolute and relative orientation tasks after VR exploration, whereas after map exploration we found the opposite pattern with lower accuracy in the pointing task (−5,7%) compared to the orientation tasks.

**Table 2:** Mean accuracies in the three spatial tasks

| Task | Absolute orientation |  | Relative orientation |  | Pointing |  |
| --- | --- | --- | --- | --- | --- | --- |
| Time | 3 seconds | infinite | 3 seconds | infinite | 3 seconds | infinite |
| VR group (%) | 50.88 | 57.2 | 49.75 | 55.43 | 52.02 | 60.61 |
| **Map group (%)** | 54.67 | 60.10 | 59.21 | 60.35 | 48.11 | 57.70 |

For the statistical analysis, as we have binary answers in our spatial tasks, we performed a logistic regression analysis with 44 participants and 9504 observations of single trials to log odds of being accurate in the time and task conditions comparing the groups that explored Seahaven in VR or with a map. We modeled the log odds of being accurate for each trial based on task (absolute, relative, pointing), time (3 seconds, infinite time condition), and exploration group (VR, map) and their interactions. The categorical variables of task, time, and group were coded using effect coding so that the log odds ratios can be directly interpreted as the main effects from the model coefficients. The model’s coefficients represent the log odds ratios of the accuracy of one categorical level from all other levels.

To test the goodness of fit of our model, we performed a log-likelihood ratio test for the full model and the null model fitted only on the intercept showing that the full model yielded a significantly better model fit (χ^2^(9) = 68.69, p<0.001).

The logistic regression model revealed a significant main effect for time (log odds ratio = 0.24, SE = 0.04, df = 9502 z = −6.03, p < 0.001), which showed that the infinite response condition resulted in overall higher accuracy levels than responding within 3 seconds. Additionally, we found a significant main effect for group (log odds ratio = −0.09, SE = 0.04, df = 9499, z = 2.33, p = 0.02) with a slightly higher accuracy by 0.09 log odds after map than VR exploration. The logistic regression showed a significant task*group interaction with a significantly higher accuracy in the pointing task in the VR group than in the map group (log odds ratio = 0.27, SE = 0.10, df = 9495, z = −2.71, p < 0.01). We found no further significant effects. All model results are displayed in Table 3.

**Table 3:** Results of the logistic regression for time*task*group comparison

| **Logistic regression (Effect coding)** | **log odds ratio** | **Standard error** | **Z-score** | **P-value** |
| --- | --- | --- | --- | --- |
| **Main effect time (Infinite)** | 0.24 | 0.04 | - 5.99 | **p < 0.001\*\*\*** |
| **Main effect task (Pointing)** | - 0.04 | 0.05 | -0.86 | p = 0.38 |
| **Main effect task (Relative)** | 0.019 | 0.05 | 0.38 | p = 0.69 |
| **Main effect group (VR)** | -0.09 | 0.04 | -2.32 | **p = 0.01\*** |
| **Task\*time interaction (Pointing, Infinite)** | 0.13 | 0.10 | 1.28 | p = 0.19 |
| **Task\*time interaction (Relative, Infinite)** | - 0.09 | 0.10 | - 0.981 | p = 0.32 |
| **Time\*group interaction (Infinite, VR)** | 0.05 | 0.08 | 0.70 | p = 0.47 |
| **Task\*group interaction (Pointing, VR)** | 0.27 | 0.10 | 2.71 | **p < 0.01\*\*** |
| **Task\*group interaction (Relative, VR)** | -0.15 | 0.10 | -1.54 | p = 0.47 |

Taken together, the spatial tasks showed a significant difference between the 3-second and infinite-time conditions with higher accuracy with time for cognitive reasoning and a significant group effect with an overall higher accuracy after map exploration. Importantly, our results revealed a significant task*group interaction with a significantly higher accuracy in the pointing task after VR exploration than after map exploration (Figure 7).

**Figure 7:**
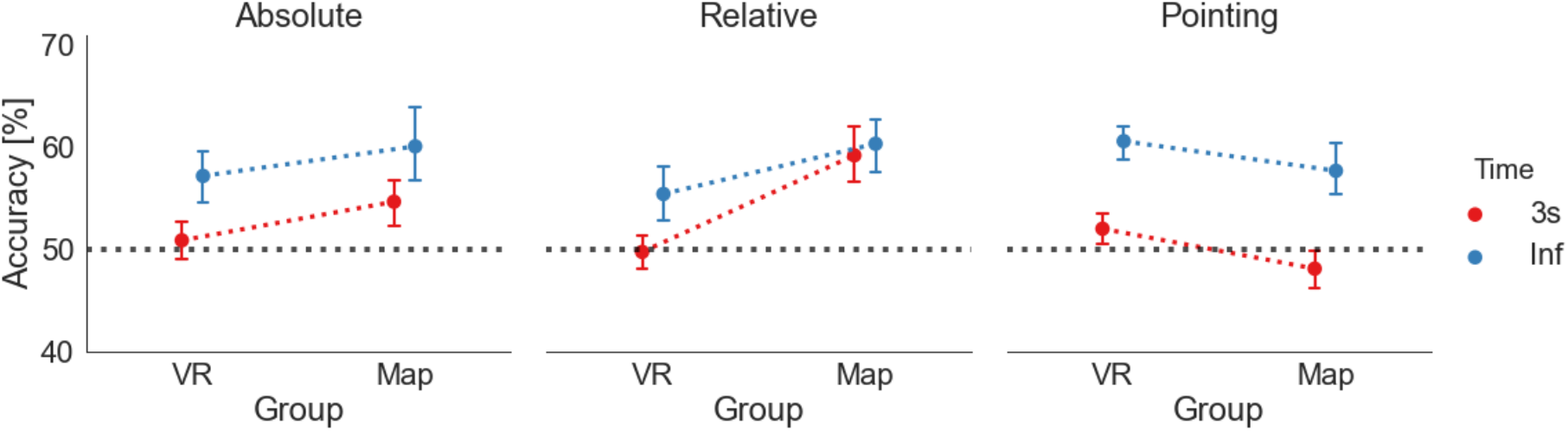
Accuracy of absolute orientation (left), relative orientation (middle), and pointing task (right) in 3 seconds (red) and infinite time (blue) conditions. In each task, the dots on the left depict the mean accuracy level after VR exploration and the dots on the right the mean accuracy level after map exploration. The dotted red (3 seconds) and blue (infinite) lines are displayed to visualize the difference between the map and VR condition. The black dashed line marks the level of 50%. The error bars indicate standard error of the mean (SEM).

### Accuracy as a Function of Familiarity of Buildings measured as Dwelling Time and Clicks on a Building

We hypothesized that buildings with a longer dwelling time would be better known and that this would increase the familiarity of the building and consequently the task accuracy. We calculated the dwelling time by counting the consecutive samples on a building recorded by the eye tracker. We considered at least seven consecutive samples on a building as one gaze event, which sums up to 233 ms as the minimum time for viewing duration. We used median absolute deviation-based outlier detection to remove outlying viewing durations. Furthermore, we defined dwelling time as the cumulative time of such gaze events spent on a particular building by a subject. In this manner, we calculated the mean dwelling time for each subject for each building. The absolute orientation task only consisted of one building in a trial, and we considered the dwelling time on this building as the dependent variable. However, as each trial in the relative orientation task consisted of a triplet of buildings, we compared the dwelling time on the prime and the two target buildings and considered the smallest value as an estimate of the minimum familiarity. Each trial of the pointing task consisted of a prime and a target building, for which we compared the dwelling time on the two buildings. In these tasks, we then conservatively used the building with the lowest dwelling time in a trial as the relevant indicator for the respective trial. With this, we calculated the average accuracy of all trials over subjects with a specific dwelling time. We calculated a Pearson correlation between accuracy and dwelling time on a building over all task and time conditions (Figure 8, left). We repeated the same analysis for comparison with the data after map exploration using the number of clicks on a building as a similar factor for familiarity with a building (Figure 8, right). After VR exploration, we found a weak significant correlation between accuracy and dwelling time (rho(216) = 0.140, p = 0.04), similar to the correlation after map exploration (rho(216) = 0.165, p = 0.02). In both exploration groups, we found a positive correlation between task accuracy and the increased familiarity of buildings measured as dwelling time on a building during VR exploration or clicks on a building during map exploration.

**Figure 8:**
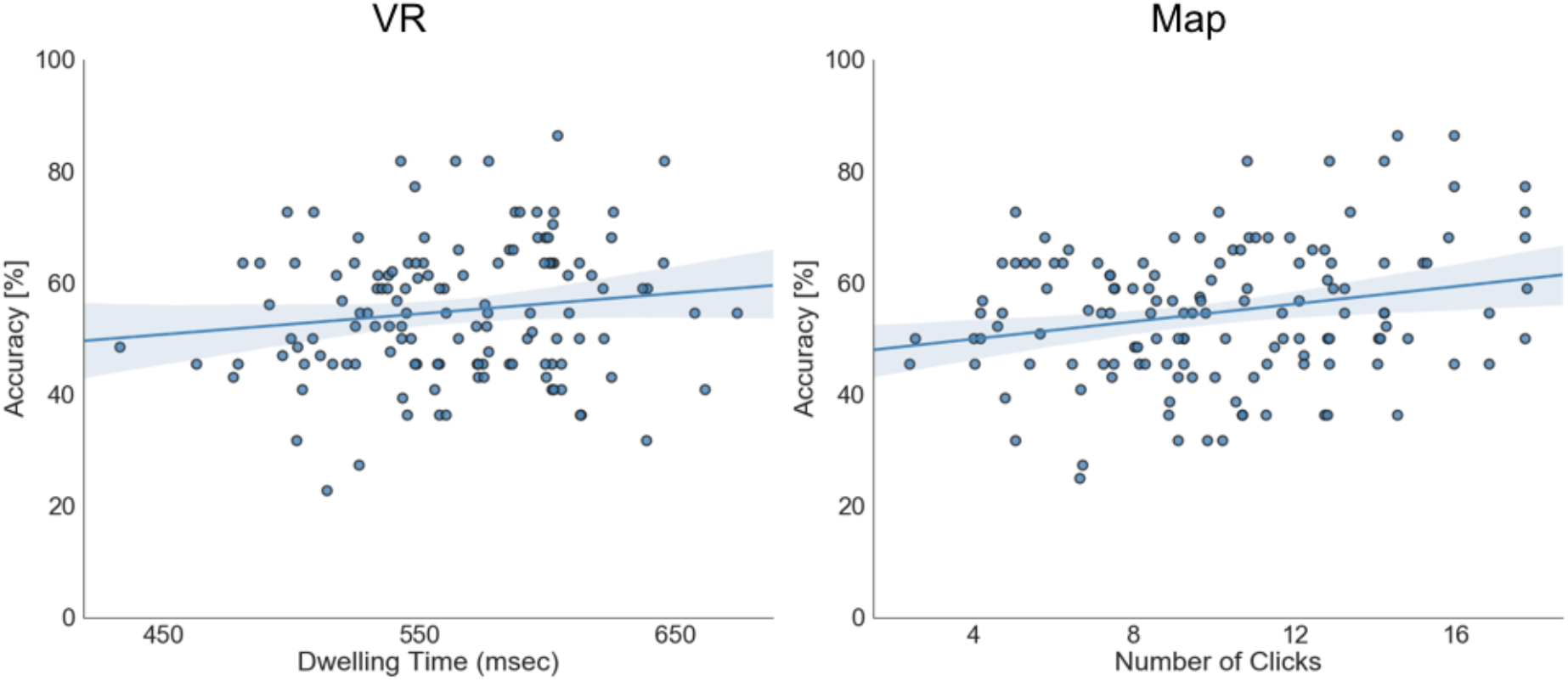
Pearson correlation between overall task accuracy and the dwelling time on a building after VR exploration (left) and the number of clicks on a building after map exploration (right). One dot represents the average accuracy of all trials over subjects with a specific dwelling time (left) or the number of clicks (right) on a building. The blue line depicts the regression line and the shaded area the 95% confidence interval.

### Accuracy as a Function of Alignment

Following previous research (e.g. König et al. 2019; Montello et al. 2004), which found a consistent alignment effect after spatial learning with a map that was less reliable after spatial learning by direct experience (e.g. Burte & Hegarty, 2014; Presson & Hazelrigg, 1984), we did not expect to find an alignment effect revealing orientation specificity after experience in the immersive virtual village opposed to map exploration. To investigate orientation specificity, we tested whether task accuracy was higher when the orientation of the building’s facing direction in the absolute orientation task was aligned with north. To obtain clear results, we only considered the absolute orientation task as participants were required to judge separate single buildings only in this task. As before, the analysis combined deviations resulting from clockwise or anti-clockwise rotations—specifically, matching bins from 0–180° and 180–360° deviations were collapsed.

To evaluate the effect of alignment to north, we used a logistic regression model with effect coding. With this, we modeled the accuracy in single trials of the absolute task given the angle (0°, 30°, 60°, 90°, 120°, 150°, and 180°), time (3 seconds or infinite condition), and exploration group (VR or map) and their interactions.

To test the goodness of fit of our model, we performed a log-likelihood ratio test to compare the full model and the null model fitted only on the intercept showing that the full model yielded a significantly better model fit (χ^2^(21) = 87.38, p<0.001).

The analysis revealed a main effect of angle at 30° (log odds ratio = 0.55, SE = 0.15, df = 3146, z = 3.61, p < 0.001) and 90° (log odds ratio = 0.39, SE = 0.15, df = 3146, z = 2.62, p < 0.01). Additionally, we found a significant angle*group interaction for 0° (log odds ratio = −0.88, SE = 0.34, df = 3146, z = −2.57, p = 0.01) and 60° (log odds ratio = −0.78, SE = 0.29, df = 3146, z = −2.66, p < 0.01) with a significantly higher accuracy for both angles after map exploration than VR exploration (Figure 9). We again confirm the significant main effect for time (log odds ratio = 0.29, SE = 0.08, df = 3146, z = 3.71, p < 0.001) and group (log odds ratio = −0.16, SE = 0.08, df = 3146, z = −2.10, p < 0.05). We did not find any further significant effects (see Table 4 for all results) (Figure 9).

**Figure 9:**
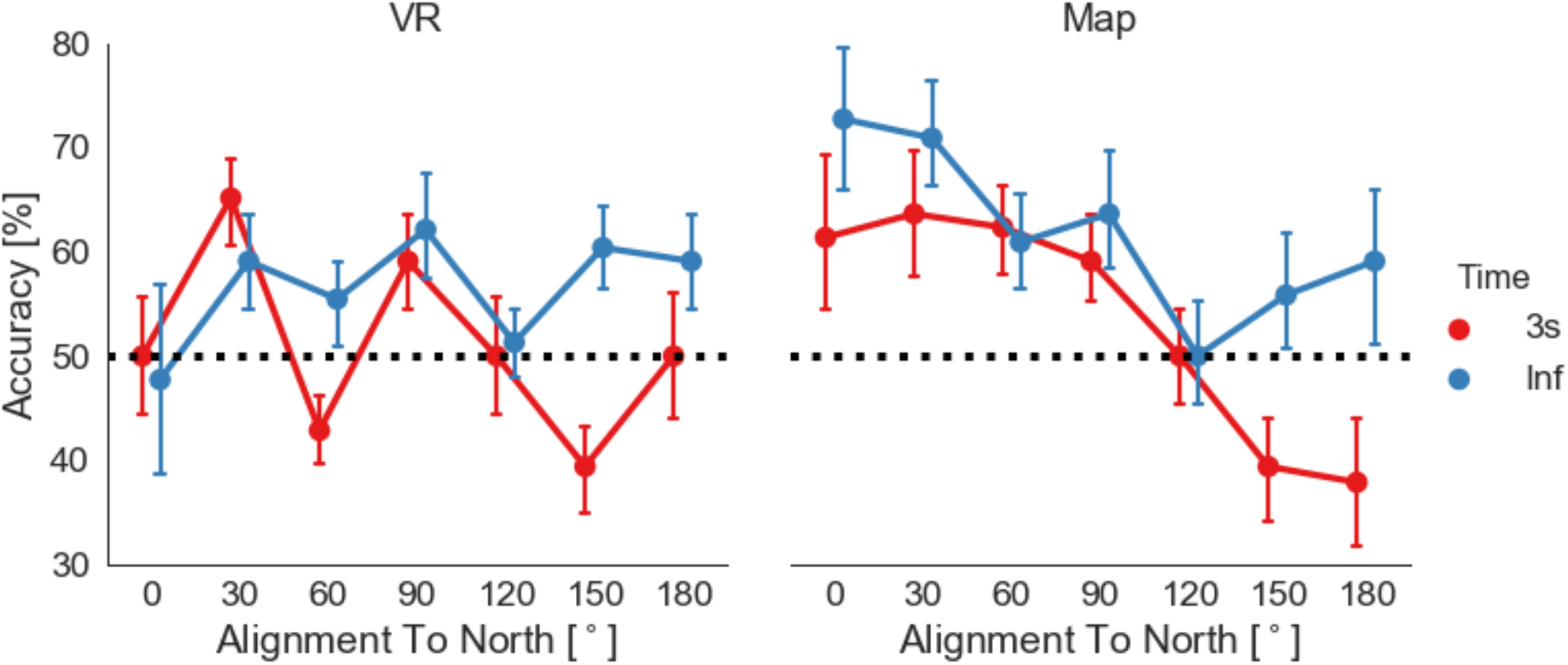
Task accuracy in the absolute orientation task in relation to the angular difference between north and tested stimuli orientation (alignment): left, after VR exploration and right, after map exploration. Dots depict mean accuracy in respect to 0°, 30°, 60°, 90°, 120°, 150°, and 180° categories. Error bars depict SEM. The black dashed line marks the level of 50%.

**Table 4:**
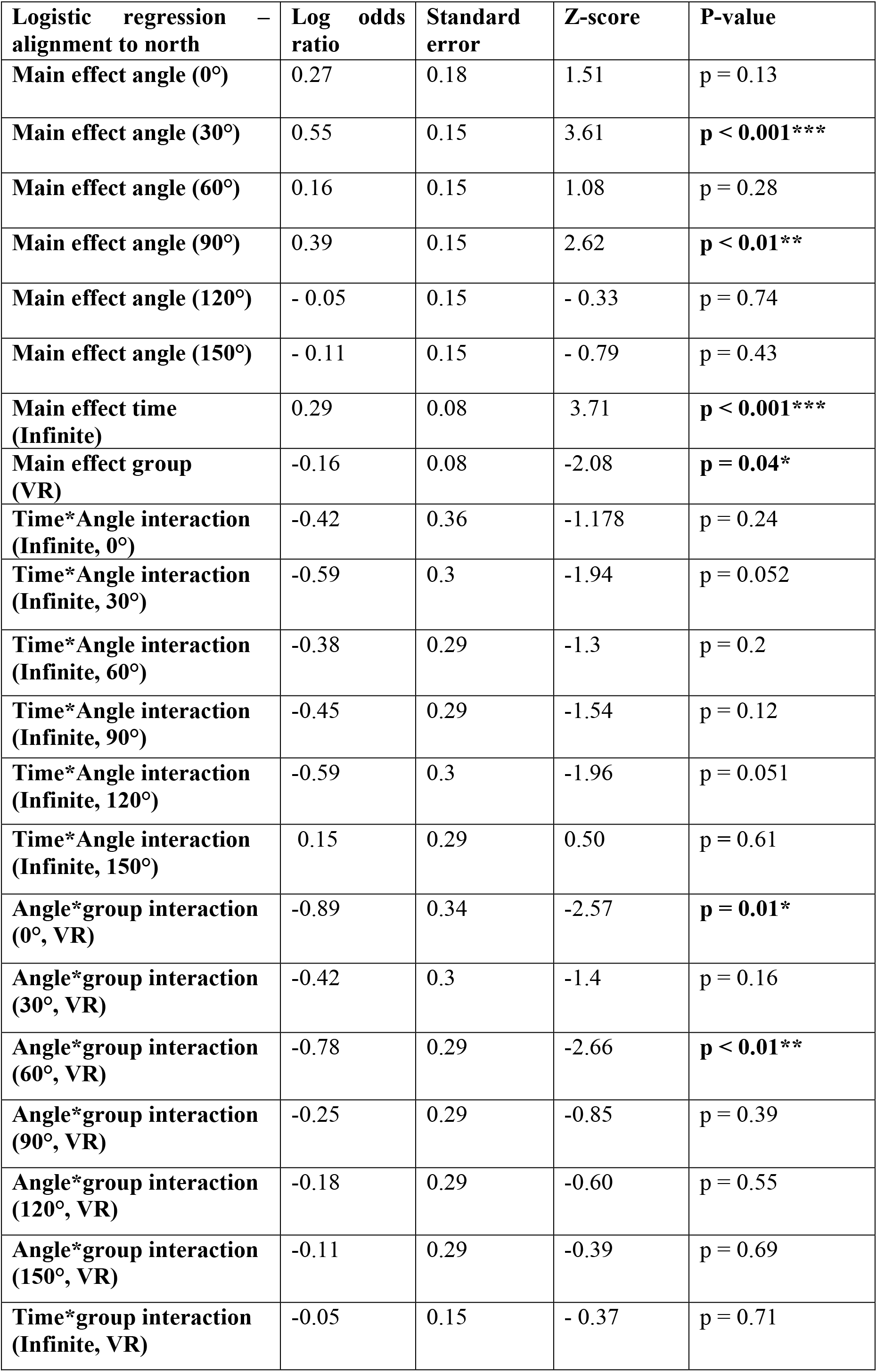
Results of the logistic regression for time*angular difference to north*group comparison

| **Logistic regression – alignment to north** | **Log odds ratio** | **Standard error** | **Z-score** | **P-value** |
| --- | --- | --- | --- | --- |
| **Main effect angle (0°)** | 0.27 | 0.18 | 1.51 | p = 0.13 |
| **Main effect angle (30°)** | 0.55 | 0.15 | 3.61 | **p < 0.001\*\*\*** |
| **Main effect angle (60°)** | 0.16 | 0.15 | 1.08 | p = 0.28 |
| **Main effect angle (90°)** | 0.39 | 0.15 | 2.62 | **p < 0.01\*\*** |
| **Main effect angle (120°)** | - 0.05 | 0.15 | - 0.33 | p = 0.74 |
| **Main effect angle (150°)** | - 0.11 | 0.15 | - 0.79 | p = 0.43 |
| **Main effect time (Infinite)** | 0.29 | 0.08 | 3.71 | **p < 0.001\*\*\*** |
| **Main effect group (VR)** | -0.16 | 0.08 | -2.08 | **p = 0.04\*** |
| **Time\*Angle interaction (Infinite, 0°)** | -0.42 | 0.36 | -1.178 | p = 0.24 |
| **Time\*Angle interaction (Infinite, 30°)** | -0.59 | 0.3 | -1.94 | p = 0.052 |
| **Time\*Angle interaction (Infinite, 60°)** | -0.38 | 0.29 | -1.3 | p = 0.2 |
| **Time\*Angle interaction (Infinite, 90°)** | -0.45 | 0.29 | -1.54 | p = 0.12 |
| **Time\*Angle interaction (Infinite, 120°)** | -0.59 | 0.3 | -1.96 | p = 0.051 |
| **Time\*Angle interaction (Infinite, 150°)** | 0.15 | 0.29 | 0.50 | p = 0.61 |
| **Angle\*group interaction (0°, VR)** | -0.89 | 0.34 | -2.57 | **p = 0.01\*** |
| **Angle\*group interaction (30°, VR)** | -0.42 | 0.3 | -1.4 | p = 0.16 |
| **Angle\*group interaction (60°, VR)** | -0.78 | 0.29 | -2.66 | **p < 0.01\*\*** |
| **Angle\*group interaction (90°, VR)** | -0.25 | 0.29 | -0.85 | p = 0.39 |
| **Angle\*group interaction (120°, VR)** | -0.18 | 0.29 | -0.60 | p = 0.55 |
| **Angle\*group interaction (150°, VR)** | -0.11 | 0.29 | -0.39 | p = 0.69 |
| **Time\*group interaction (Infinite, VR)** | -0.05 | 0.15 | - 0.37 | p = 0.71 |

Our results revealed a significant main effect for 30° and 90° angular differences to north. Furthermore, they showed that 0° and 60° differences to north had significantly higher accuracies after map exploration than after VR exploration, thus suggesting an alignment effect only after spatial learning with a map.

As one reason for the lack of an alignment effect toward north after VR exploration, we assumed that participants did not know where the defined north was in the virtual village. To investigate how well participants deduced the cardinal directions from the virtual sun’s position in VR, we asked them at the end of each exploration to turn so that they would face their subjectively estimated north (Figure 10, left). Comparing participants’ subjective estimations of north after the third exploration of Seahaven with true north in the virtual village revealed that, overall, participants’ knowledge of north was remarkably accurate. The estimations of 16 out of 22 participants lay within 45° deviation from true north and six below 3°. Nevertheless, we found no correlation between deviation from true north and accuracy (Pearson correlation between the accuracy in the absolute orientation task and deviation from north with 3 seconds condition: rho(22) = 0.065, p = 0.78 and with infinite time condition: rho(22) = −0.02, p = 0.94). Note that we collapsed the full circle to angles from 0 to 180° for all analyses (Figure 10, right). Therefore, our results suggest that in spite of having a relatively accurate estimation of the northerly direction in the virtual village, participants could not use it to increase their knowledge of the orientation of buildings in Seahaven.

**Figure 10:**
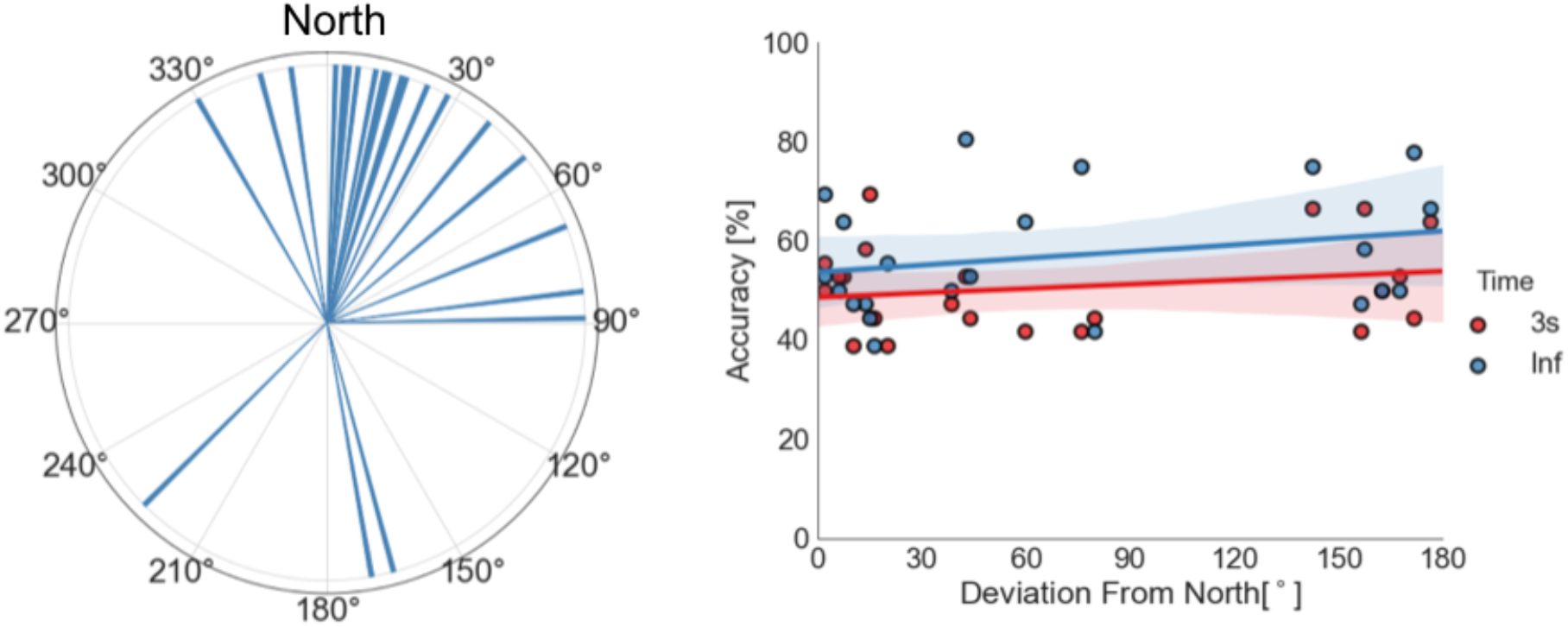
Left: Polar histogram with a bin size of 1° showing the subjective estimation of north after VR exploration by the participants. Up is true north. Right: Correlation between accuracy in the absolute orientation task and deviation from north for the 3-second response condition in red and infinite-response condition in blue. One dot depicts the combination of accuracy and deviation from the north for one participant. Lines depict regression lines (3-second response condition in red and infinite-response condition in blue). Colored shaded areas depict the respective 95% confidence intervals.

### Accuracy as a Function of Angular Difference of Choice Options

We hypothesized that accuracy would improve with larger angular differences between choice options of our 2AFC tasks independent of whether exploration was in the virtual environment or using the map. Participants in both exploration groups performed the same spatial tasks. Thus, all participants were required to choose between two alternative choices that differed from each other in varying angular degrees in steps of 30°. The analysis combined deviations resulting from clockwise or anti-clockwise rotations. Therefore, matching bins from 0–180° and 180–360° deviations were collapsed. Due to small variations in the orientation of the buildings in the virtual village design, the difference between these angles varied in the relative orientation task with a maximal deviation of +/− 5° in each step from a minimum of 30° in steps of 30°. This design resulted for all tasks in bins of 30°, 60°, 90°, 120°, 150°, and 180° angular difference. For each bin, the accuracy was calculated combining all tasks but separately for the 3-second and infinite-time conditions.

For statistical analysis, we performed another logistic regression (Table 5) to model the log odds ratio of being accurate based on angle (the angular difference between decision choices: either 30°, 60°, 90°, 120°, 150°, or 180°) and time (3 seconds or infinite time condition) and exploration group (VR or map) (Figure 11). As discussed above, the categorical variables were coded with effect coding and hence the model coefficients can be directly interpreted as main effects.

**Figure 11:**
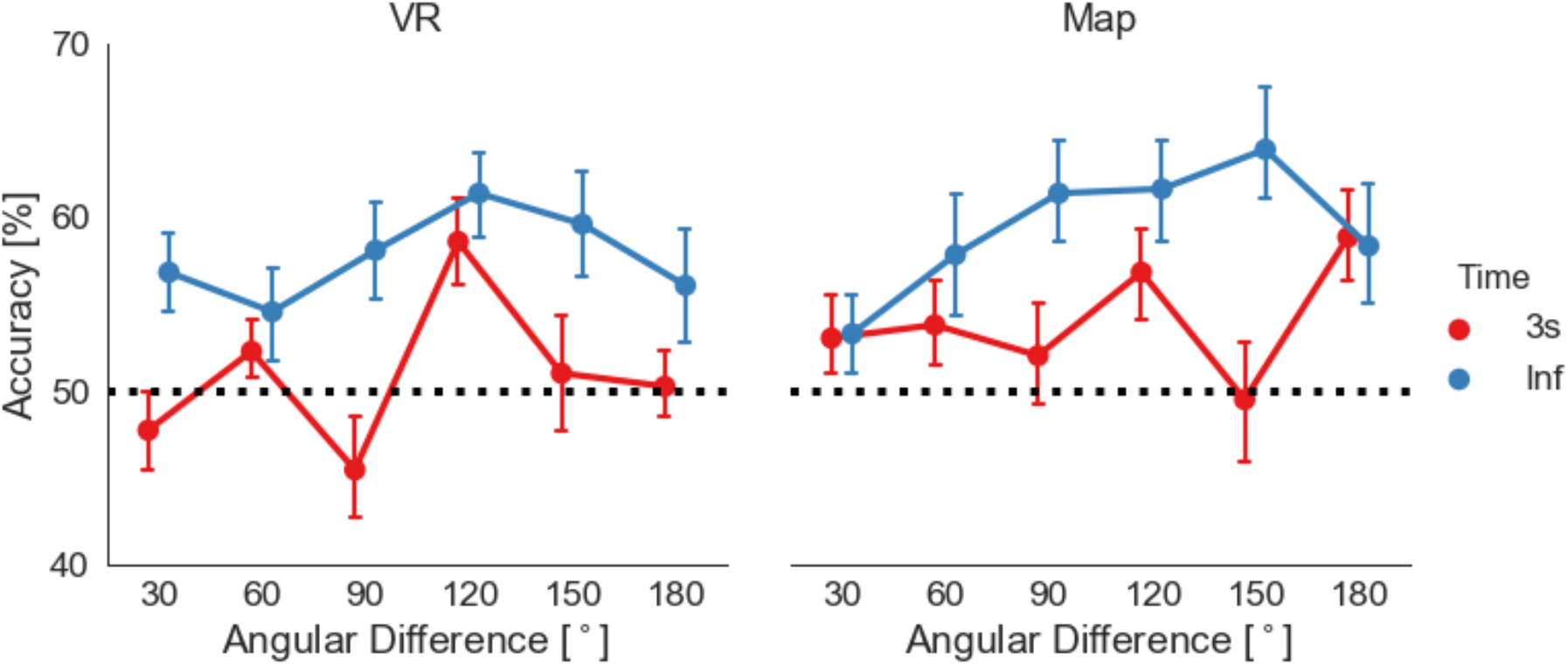
Overall task accuracy in relation to the angular difference between choices in the task stimuli after VR exploration (left) and map exploration (right). Dots depict mean accuracy in respect to 30°, 60°, 90°, 120°, 150°, and 180° categories, with a response within 3 seconds in red and with unlimited time in blue. Error bars represent SEM. The black dashed line marks the level of 50%.

**Table 5:** Results of the logistic regression for time*angular difference*group comparison

| Logistic regression – angular difference | log odds ratio | Standard error | Z-score | P-value |
| --- | --- | --- | --- | --- |
| Main effect angle (60°) | 0.07 | 0.07 | 1.06 | $p = 0.28$ |
| Main effect angle (90°) | 0.06 | 0.07 | 0.88 | $p = 0.37$ |
| Main effect angle (120°) | 0.28 | 0.07 | 3.90 | **$p < 0.001^{***}$** |
| Main effect angle (150°) | 0.13 | 0.07 | 1.89 | $p = 0.06$ |
| Main effect angle (180°) | 0.12 | 0.07 | 1.78 | $p = 0.07$ |
| Main effect time (Infinite) | 0.25 | 0.04 | - 6.00 | **$p < 0.001^{***}$** |
| Main effect group (VR) | -0.09 | 0.04 | -2.32 | **$p = 0.02^*$** |
| Time*Angle interaction (Infinite, 60°) | -0.06 | 0.14 | - 0.42 | $p = 0.67$ |
| Time*Angle interaction (Infinite, 90°) | 0.25 | 0.14 | 1.80 | $p = 0.07$ |
| Time*Angle interaction (Infinite, 120°) | -0.03 | 0.14 | - 0.20 | $p = 0.83$ |
| Time*Angle interaction (Infinite, 150°) | 0.28 | 0.14 | 1.96 | **$p = 0.05^*$** |
| Time*Angle interaction (Infinite, 180°) | -0.08 | 0.14 | -0.56 | $p = 0.57$ |
| Angle*group interaction (60°, VR) | -0.06 | 0.14 | -0.43 | $p = 0.66$ |
| Angle*group interaction (90°, VR) | -0.16 | 0.14 | -1.15 | $p = 0.24$ |
| Angle*group interaction (120°, VR) | 0.06 | 0.14 | 0.46 | $p = 0.63$ |
| Angle*group interaction (150°, VR) | -0.02 | 0.14 | -0.14 | $p = 0.88$ |
| Angle*group interaction (180°, VR) | -0.18 | 0.14 | -1.29 | $p = 0.19$ |
| Time*group interaction (Infinite, VR) | 0.05 | 0.08 | 0.69 | $p = 0.48$ |

To test the goodness of fit of our model, we performed a log-likelihood ratio test for the full model and the null model fitted only on the intercept showing that the full model yielded a significantly better model fit (χ^2^(18) = 77.29, p<0.001).

Our logistic regression analysis showed a main effect for time and for group with a significant higher accuracy for the infinite time condition (log odds ratio = 0.25, SE = 0.04, df = 9502, z = −6.0, p < 0.001) and a higher accuracy in the map group (log odds ratio = −0.09, SE = 0.07, df = 9496, z = 2.32, p = 0.02). The analysis also revealed a significantly higher accuracy for 120° angular difference between task choices compared to all other angular differences (log odds ratio = 0.28, SE = 0.07, df = 9497, z = 3.90, p <0.001). Additionally, we found a significant time*angle interaction with a significantly higher accuracy in the infinite time condition for 150° angular difference between choice options (log odds ratio = 0.28, SE = 0.14, df = 9491, z = 1.96, p = 0.05). We found no further significant effects (see Table 5 for all results) (Figure 11).

To summarize, analyzing the effect of angular difference of choice options on task accuracy, we found a significant main effect of angle, independent of the source of exploration, with higher accuracy with 120° angular difference between task choices. Additionally, we found a significantly higher accuracy for 150° difference between task choices for the infinite response time condition than for the 3-second time condition.

### Accuracy as a Function of Distance

In line with previous research (Meilinger, Frankenstein, Watanabe, Bülthoff, & Hölscher, 2015), we hypothesized a higher task accuracy with smaller distances between tested buildings (distance effect) with experience in the immersive virtual environment. As we have to be able to compare the distance between two buildings for this analysis, we only considered the relative orientation and the pointing task. In the pointing task, we had only two buildings in a trial: the prime and the target, whose distance we directly compared. In the relative orientation task, we compared the distance between the prime and the correct target building only, leaving out the wrong target building for this analysis. So, distance is defined as the distance between predefined building pairs. For the analysis, we averaged the distance of building pairs over subjects.

To investigate the dependencies of accuracy on the distance, we calculated an ordinary least squares linear regression model with accuracy as the dependent variable (Figure 12). Here, we focus on the effects of distance and the modulation by time and group as the independent factors. Our linear regression model for the relative orientation task revealed (Table 6 and Figure 12, left) a main effect for distance (beta = −0.04, t(137) = −2.89, p = 0.004). This showed that, with a one-unit change of distance, accuracy decreased by 0.04%. Furthermore, we found a two-way interaction for time*distance (beta = 0.08, t(137) = 2.59, p = 0.01), which shows that time conditions affected distance differentially with the 3-second response condition resulting in a larger decrease in accuracy (by 0.08% for one unit change in distance) in comparison to the infinite time condition. We did not find any further significant results for the relative or pointing tasks (Table 6 and Figure 12).

**Figure 12:**
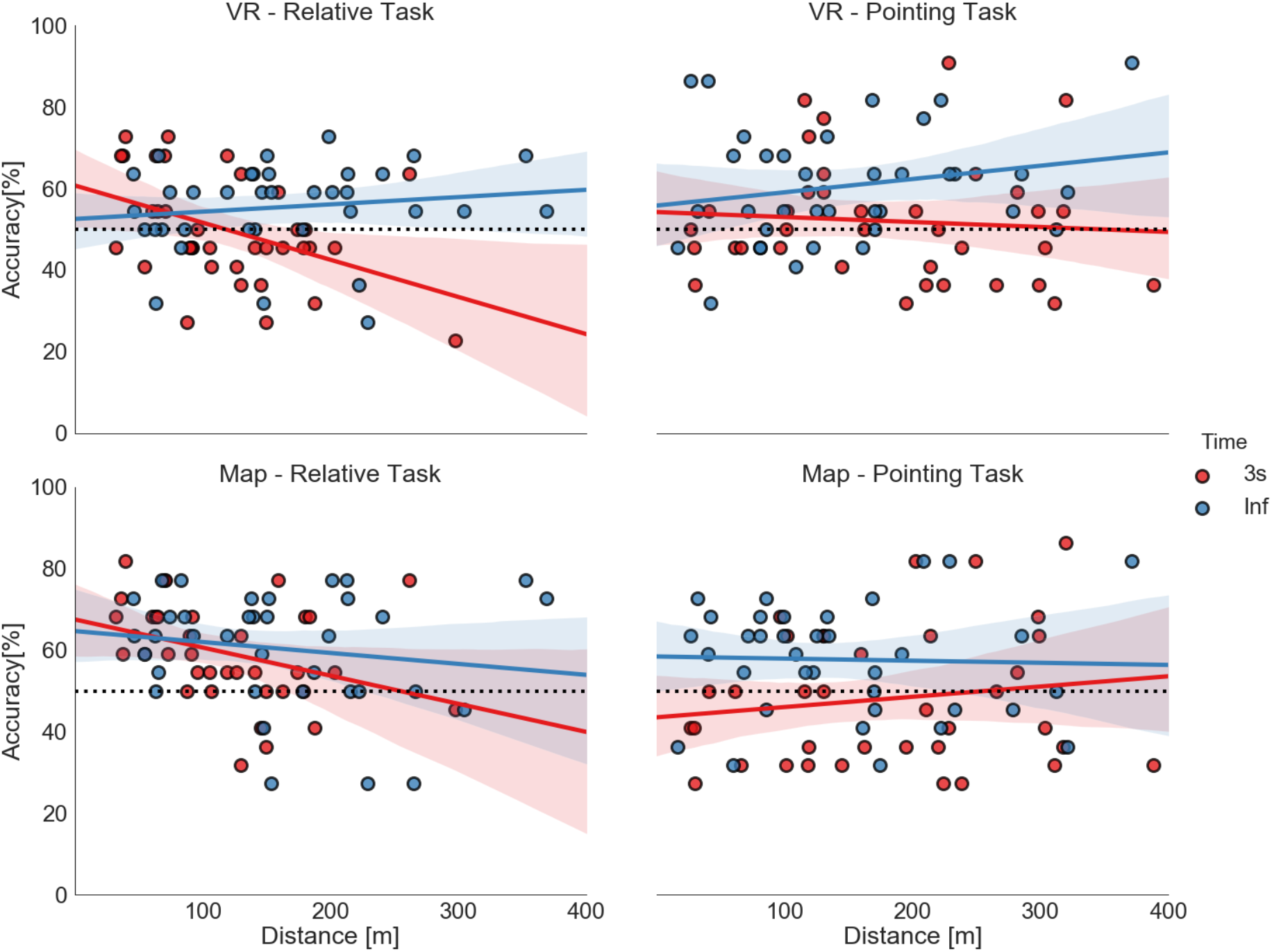
Correlation of distance (abscissa) and task accuracy (ordinate) (top—VR; bottom—map). The dots depict the combination of time and distance averaged over subjects for a building pair: 3-second response condition in red and infinite-response condition in blue, on the left side for the relative orientation task and on the right side for the pointing task. The straight lines depict the correlation lines and the colored shaded areas the respective 95% confidence interval. The black dashed line marks the level of 50%.

**Table 6:** Results of the linear regression model for distance

| Linear regression for distance – relative orientation task | Beta | T-value | P-value |
| --- | --- | --- | --- |
| Main effect distance | beta = - 0.04 | t(137) = - 2.89 | p < 0.01** |
| **Time\*distance interaction**<br>**(3 seconds, distance)** | beta = 0.08 | t(137) = 2.59 | **p = 0.01\*** |
| **Group\*distance interaction**<br>**(map, distance)** | beta = 0.02 | t(137) = 0.68 | p = 0.5 |
| **Group\*time\*distance**<br>**interaction**<br>**(map, 3 seconds, distance)** | beta = 0.06 | t(137) = 1.07 | p = 0.29 |

| **Linear regression for distance –**<br>**pointing task** | **Beta** | **T-value** | **P-value** |
| --- | --- | --- | --- |
| **Main effect distance** | beta = 0.01 | t(137) = 0.75 | p = 0.46 |
| **Time\*distance interaction**<br>**(3 seconds, distance)** | beta = 0.007 | t(137) = 0.27 | p = 0.79 |
| **Group\*distance interaction**<br>**(map, distance)** | beta = 0.001 | t(137) = 0.04 | p = 0.97 |
| **Group\*time\*distance**<br>**interaction**<br>**(map, 3 seconds, distance)** | beta = 0.08 | t(137) = 1.4 | p = 0.16 |

Taken together, these results unexpectedly showed the same influence pattern of distance after exploration in VR and with a map. In both groups, a distance effect was only visible with higher accuracy for shorter distances between tested buildings in the relative orientation task with a response within 3 seconds.

### Accuracy as a Function of FRS Scaling

To estimate subjectively rated abilities in spatial-orientation strategies learned in real-world environments, we asked subjects to fill out the FRS questionnaire (Münzer et al., 2016b; Münzer & Hölscher, 2011). To investigate the influence of these spatial-orientation abilities on task accuracies after exploring a virtual village, we performed the more robust Spearman’s rank correlation analysis of the FRS scales and task accuracies and reanalyzed the map data accordingly. After VR exploration, we did not find significant correlations with any of the task and time accuracies (p values between 0.09 and 0.91) for scale 1 (the global-egocentric orientation scale). Similarly, on scale 2, the survey scale, the results showed no significant correlation (p values between 0.06 and 0.46). For scale 3, the cardinal directions scale, we found a correlation with the absolute orientation task with unlimited response time (rho(19) = 0.51, p = 0.03). No other correlations for scale 3 were significant (p values between 0.13 and 0.98). After map exploration, we found a correlation between Scale 1 and the pointing task with a 3-second response time (rho(19) = 0.488, p = 0.03). No other correlations were significant (p values between 0.07 and 0.87). As the p values did not survive Bonferroni multiple comparison correction (p < 0.002), we are cautious about interpreting these results any further.

## Discussion

In the present study, our research question was whether and how spatial learning in a virtual environment is influenced by different sources of spatial knowledge acquisition, comparing experience in the immersive virtual environment with learning of an interactive city map of the same environment (König et al., 2019). Taken together, our results showed a significantly higher accuracy when unlimited time was given for cognitive reasoning than for a response within 3 seconds, independent of the source of exploration and with a slight overall higher accuracy after map exploration. Furthermore, we found a higher accuracy for judging straightline directions between buildings after exploration in the immersive VR than after exploration with the map. After map exploration, we found a higher accuracy of cardinal directions and building-to-building orientation tasks. Increased familiarity of buildings was weakly correlated with an increase in task accuracy after VR and map exploration. We found an alignment-to-north effect only after map exploration; in contrast to exploration in immersive VR, where we found no orientation dependency of task accuracies. Our results revealed the same pattern for distances after VR and map exploration, with a distance effect visible with higher accuracy for shorter distances only for judging relative orientations of buildings within 3 seconds. We found no significant correlation of task accuracies and the scales of the FRS questionnaire in either group. Overall, our results suggest that spatial knowledge acquisition in a virtual environment was influenced by the source of spatial exploration with higher accuracy for action-relevant judgment of straight-line directions between buildings after exploration in immersive VR and superior knowledge of cardinal directions and building-to-building orientation after map exploration.

When comparing the exploration of a virtual environment with real-world navigation, an important question is the extent to which these two scenarios are comparable. More specifically, we ask whether experience in a virtual environment can be compared to direct experience in the real world. For experience in VR to be comparable to direct experience in the real world it has to provide sensory input that is as natural as possible. Therefore, we used an immersive virtual environment setup with an HMD. Participants had the feeling of being surrounded by the virtual environment with a 360° panoramic scenery, which supported the feeling of presence in the VR. Furthermore, the HTC Vive headset that we used provided a field-of-view of 110° that also allowed peripheral vision, which is considered to be important for developing survey knowledge (Sholl, 1996). For the experience of movement, we provided natural visual flow and physical-body rotation on a swivel chair combined with forward movement that was controlled by the participants with a handheld controller. This setup was shown by Riecke et al. (2010) to result in comparable performance levels to those of full walking. With this design, our VR environment might be considered as a primary environment that could be directly experienced in a manner more similar to the direct experience of a natural environment compared to indirect sources like maps (Montello et al., 2004; Presson & Hazelrigg, 1984). Our interactive city map provided participants with a north-up city map from a bird’s eye perspective and additionally screenshots of the buildings’ front-views from a pedestrian perspective, thus providing a feature that is not typical for regular city maps. Even though, we assume that our interactive city map is more like an indirect source than direct experience for spatial learning. Thus, although direct experience in the real world differs in many aspects from the experience in virtual environments, we suggest that Seahaven is a valid virtual environment to investigate spatial learning by experience in immersive VR and allows the comparison to spatial learning with the use of an interactive map of the same VR environment as an indirect source.

For the design of our virtual village, we considered several important points. Investigating real-world large-scale environments poses the problem of controlling performed spatial learning. This is solved in some studies by investigating a natural environment of a manageable size, such as floors in a building (Richardson, Montello, & Hegarty, 1999b; Thorndyke & Hayes-Roth, 1982) or University campuses (Burte & Hegarty, 2012; Montello et al., 2004; Sholl et al., 2006). In comparison to these real-world environments, Seahaven has a substantial size. The structure of the environment also influences spatial learning. The grid-like structure of, for example, Manhattan and other American cities allows spatial navigation to progress by counting the number of street crossings straight ahead and to the right and left for orientation purposes. In contrast, we used an unordered street plan for our virtual village as observed in most European countries to avoid this kind of navigation and to present a VR environment matching the style of the city the participants lived in. Taken together, our virtual environment is tuned to support spatial learning similarly to spatial knowledge acquisition under natural conditions.

The choice of tasks in the present study was motivated by previous research investigating the influence of the feelSpace sensory augmentation device, which gives information of cardinal north via vibrotactile information on space perception (Kaspar et al., 2014; König et al., 2016). In these studies, participants who trained with the feelSpace device reported various alignment effects in relation to allocentric reference cues and inter-object relations. To investigate these reported effects behaviorally (König et al., 2017), we designed three spatial tasks to investigate the learning of allocentric knowledge in relation to the cardinal directions, the relative orientations between objects, and judgment of relative directions between locations in the environment. In this previous study, we used photographs of buildings and streets of a real-world city as stimuli as buildings and streets are important landmarks for spatial navigation in real-world cities. In the presented study, we also focused on buildings as our objects of interest and performed the same spatial-navigation tasks with screenshots of buildings of the virtual village after VR and map exploration.

In detail, the absolute orientation task evaluates knowledge of single buildings’ orientation in respect to cardinal directions. The relative-orientation task evaluates the knowledge whether two buildings share the same orientation, thus testing building-to-building orientation. This task can be solved in different ways. One way would be to memorize the cardinal orientation of single buildings, which will then be compared with each other to deduce the relative orientation between two buildings. Another way would be to memorize whether two buildings are aligned within the city layout directly memorizing the relative orientation of a pair of buildings. This way would not require knowledge of single building’s orientation towards north. In fact, a previous study (König et al., 2017) argues that some part of spatial information is encoded in the former, a different part in the latter fashion. As such, knowledge of cardinal directions might be used to solve this task, but are not necessary to do so. Thus, the relative orientation task is designed to test knowledge of the relative orientation of two buildings without reference to their absolute orientation, e.g. north, or to their location and tests a non-redundant complementary aspect of spatial knowledge in the present set of tasks. The pointing task evaluates judgment of the direction from the location of one building to the location of another building. The task was designed as a straight-line pointing task from one building to another building in close analogy to the “orientation task” of Thorndyke and Hayes-Roth, (1982). There, participants had to indicate the direction from a start location (a room in a building complex) to a destination location (another room). Even though Thorndyke and Hayes-Roth termed their task orientation task, equally to our task the straight-line direction between locations had to be judged. Taken together, we used spatial tasks that evaluate spatial knowledge based on different allocentric reference cues.

As the sources of information for spatial learning provide different information, the learned and memorized spatial knowledge is also expected to differ. Investigating spatial knowledge acquisition after direct experience in real-world surroundings, previous research revealed that landmark, route, and survey knowledge may evolve (Montello, 1998; Richardson et al., 1999; Siegel & White, 1975; Thorndyke & Hayes-Roth, 1982), even though there are large individual differences (Hegarty, Burte, & Boone, 2018; Ishikawa & Montello, 2006; Montello, 1998; Wiener, Büchner, & Hölscher, 2009). Direct experience supports route knowledge in relation to an egocentric reference frame whereas spatial learning with a map heightens survey knowledge in relation to an allocentric reference frame (Frankenstein, Mohler, Bülthoff, & Meilinger, 2012; Meilinger et al., 2013; Meilinger et al., 2015; Montello et al., 2004; Richardson et al., 1999; Shelton & McNamara, 2004; Taylor et al., 1999; Thorndyke & Hayes-Roth, 1982). As a map directly provides a fixed coordinate system and the topography of the environment, survey knowledge is supposed to be preferentially derived from maps (Montello et al., 2004; Thorndyke & Hayes-Roth, 1982). But it can also be obtained from direct navigation within an environment (Klatzky et al., 1990; Montello, 1998; Siegel & White, 1975; Thorndyke & Hayes-Roth, 1982). To clarify spatial learning from different sources, Thorndyke and Hayes-Roth, (1982) investigated acquired spatial properties of a complex real-world building after navigation therein and compared it to map use. Their results revealed that after map learning, participants were more accurate in their estimation of Euclidian distances between locations and location of objects in the tested environment. After navigation experience, participants were more accurate at judging route distances and relative directions from a start to a destination location, which is considered to test survey knowledge (Montello et al., 2004). In the presented study, we performed a judgment-of-direction task between the location of buildings, similar to Thorndyke and Hayes-Roth, (1982). In line with Thorndyke and Hayes-Roth, (1982), we found significantly higher accuracy in the judgment of direction task after direct exploration of the virtual environment than after map exploration. Thorndyke and Hayes-Roth, (1982) argued that participants using a map were required to perform a perspective switch, from the bird’s-eye perspective of the map to the required ego perspective in the judgment of direction task. They assumed that this perspective switch might have been the cause for the lower performance after map learning. Our participants who explored the interactive map had to perform a switch of perspective even during the exploration between bird’s-eye perspective of the map to the required ego perspective in viewing the screenshots of the buildings, which was the same perspective in the judgement of direction task. Participants who explored the immersive VR used the ego perspective equally during exploration and testing. Thus, the argument that the switch of perspective might be the reason for the lower accuracy after map learning holds true also for our results. More recent research supported the view that a switch in required perspective caused a reduction in performance (McNamara, Sluzenski, & Rump, 2008; Meilinger et al., 2013; Shelton & McNamara, 2004). However, in accordance with Thorndyke and Hayes-Roth, (1982), who assumed that survey knowledge could also be achieved with enough familiarity of the environment by navigation experience, we suggest that our participants acquired some kind of spatial topographical overview after exploration of our immersive virtual environment. In contrast our tasks that evaluated knowledge of cardinal directions and relative orientations between buildings revealed higher accuracies after map exploration, even though they had to perform the same perspective switch as in the pointing task. We assume that as a map directly provides the information of cardinal directions and relative orientations, it was easier to learn this spatial information during map exploration than exploration of the immersive VR. In summary, our results suggest that judgment of straight-line directions between locations was preferentially acquired after exploration of the immersive VR in contrast to tasks using information about cardinal directions and building-to-building orientation that was preferentially derived with an interactive map.

The kind of spatial knowledge that is acquired likely also depends on the particular task (Meilinger et al., 2013). In spatial navigation, actively navigating from one place to another preferably using the shortest or fastest way without getting lost is an important daily challenge. Humans use direct experience and indirect sources such as maps to successfully navigate an environment, thus combining multiple types of knowledge. Previous research investigated spatial knowledge of the city of Osnabrück after at least one year of living in the city (König et al., 2017). Thus, in that study, direct experience and cartographic information were likely combined in acquiring knowledge of the environment. The researchers performed the same spatial tasks as in the presented study, evaluating photographs of real-world buildings of the city of Osnabrück (König et al., 2017). They found the highest accuracy in the pointing task in which participants judged relative directions of locations and the lowest accuracy for the knowledge of cardinal directions. In the present study after experience in the immersive VR, accuracy for judgment of relative directions of locations was higher than for knowledge of cardinal directions with the reverse knowledge pattern after map learning. Thus, our results after experience in immersive VR resemble the results after knowledge acquisition in the real world more than those after map learning. König et al. (2017) argued that, in line with theories of embodied cognition (Engel et al., 2013; O’Regan & Noe, 2001; Wilson, 2002), knowledge of directions between locations that enables direct action to navigate from one place to another is an important reason for spatial learning. Therefore, action-relevant information provided by experience in an immersive virtual environment or real environment might support learning of the judgment of directions between locations.

In addition to the source of spatial exploration and the performed task, the design of the task also has to be considered as an influential factor in spatial knowledge acquisition (Montello et al., 2004). Pointing tasks in spatial research generally test judgment of directions from a specific location to another specified location. Besides this general structure, there exist a large variety of different designs (e.g. McNamara, 2002; McNamara, Rump, & Werner, 2003; Montello et al., 2004; Mou & McNamara, 2002; Shelton & McNamara, 1997; Waller & Hodgson, 2006). In recent research, Zhang et al. (2014) investigated the development of cognitive maps after map and desktop VR learning with two different pointing tasks adapted from previous research (Holmes & Sholl, 2005; Mou, McNamara, Valiquette, & Rump, 2004; Waller & Hodgson, 2006). Zhang et al. (2014) used a scene and orientation-dependent pointing (SOP) task, which provided egocentric embodied scene and orientation information, and a judgment of relative direction (JRD) task, which did not give any orientation information and was presumed to be based on remembered knowledge of allocentric relations between landmarks. After spatial exploration in an immersive VR, our pointing task was performed outside the VR and displayed screenshots of the tested buildings in a 2AFC design. With this design, we stayed within the visual domain for testing. This allowed the use of identical test conditions after VR and map learning. Zhang et al. (2014) found a superior performance in the SOP task after spatial learning in VR. In contrast, after map learning of the VR environment, the performance was superior in the JRD task in that study. We found a higher accuracy with our pointing task after exploration in the immersive VR, which is in agreement with the results of the SOP task of Zhang et al. (2014). We suggest that even though the pointing stimuli did not provide scene and orientation information further to the screenshots taken in VR, the display of these visual images triggered the remembered visual scene information that was available while exploring the virtual environment. As the map exploration solely provided the screenshots of the buildings and the city map from a bird’s eye view, no additional embodied information was learned. Our results suggest that the accuracy in our pointing task was supported by remembered scene and orientation information after exploration in VR, which led to higher accuracy than after map exploration.

In everyday navigation, spontaneous decisions and careful deductive planning are combined. This is in line with dual-process theories (Evans, 1984; Evans, 2008; Finuncane, Alhakami, Slovic, & Johnson, 2000; Kahneman & Frederick, 2002), which distinguish rapid, intuitive cognitive processes, “System 1” processes, from slow, deductive, analytical cognitive processes, “System 2” processes. Kahneman & Frederick (2002) suggested that “System 2” processes, once they achieved greater proficiency, descend to and improve “System 1” processes. Thus, to investigate which kind of cognitive processes were used to solve our tasks, we investigated two response time conditions in the present study. We used a response window that required a response within 3 seconds, testing a rapid decision, and an infinite time condition with unlimited time for a response, allowing for analytical cognitive reasoning. Our results revealed a significantly higher accuracy with unlimited time to respond that allowed for slow, deductive cognitive reasoning than responses within 3 seconds after VR as well as map exploration. Thus, our results suggest that System 2 deductive cognitive processes contributed to solve our spatial tasks after VR and map exploration of Seahaven.

In real-world spatial navigation studies, orientation specificity and distance between tested objects were found to be differential factors after direct experience in the environment and map use. Orientation specificity is measured as an alignment effect that is visible in higher accuracy when learned and retrieved orientations were aligned. This was consistently found after spatial learning that provided a fixed reference frame such as cardinal directions of a map or environmental features (Brunyé, Burte, Houck, & Taylor, 2015; Frankenstein et al., 2012; McNamara, 2003; McNamara et al., 2003; Montello et al., 2004). After direct experience in the environment, which might lead to multiple local reference frames (Meilinger, 2008; Meilinger et al., 2015, 2006; Montello et al., 2004) an alignment effect was much less consistently found (Burte & Hegarty, 2014; Presson & Hazelrigg, 1984). In agreement, our results revealed no alignment effect of tested building orientations toward north after experience in the immersive virtual village. Even though participants were remarkably accurate at estimating north after VR exploration, they were not able to use this knowledge to improve task accuracies. In contrast, we found an alignment effect toward north after map exploration (König et al., 2019). Thus, our results are in line with a differential effect of orientation specificity after experience in immersive VR and map exploration.

Investigating the influence of the distance between tested buildings, previous research in real-world environments indicated that when acquiring knowledge of a large-scale environment by direct experience, smaller distances between tested objects improved task performance (distance effect), whereas after learning with a map no distance effect was found (Frankenstein et al., 2012; Loomis et al., 1993; Meilinger et al., 2015). In contrast, our results revealed the same pattern of distance dependency after village exploration in VR and with a map. In both groups, with time for cognitive reasoning to respond, task accuracies were not influenced by the distance between tested buildings whereas when responses within 3 seconds were required, we found a significant negative correlation between accuracy and distance in judging the relative orientation between buildings but not in judging the direction between buildings locations in the pointing task. The spatial extent of our virtual village, which allowed for distances between buildings up to 400 (virtual) meters, is relatively large for a virtual environment. However, compared to a real-world city, this distance is small and might not be large enough to yield distance effects as in a larger real-world city. Thus, our results revealed a comparable pattern for the influence of distance on task accuracy after VR and map exploration with only a distance effect in judging building-to-building orientation with responses within 3 seconds, but the extent of our virtual village might not be large enough to reliably investigate this effect.

Real-world navigation is a multimodal activity that requires embodied interaction of the navigator with the environment being in line with theories of embodied and enacted cognition, which understand cognition as an embodied activity that includes mind, body, and environment (Engel et al., 2013; Noë, 2004; O’Regan & Noë, 2001; Varela, Thompson, & Rosch, 1991; Wilson, 2002). Spatial navigation continuously requires multimodal sensory input about visual, motor, kinesthetic, and vestibular changes. Thus, gaining more insight into natural navigation with the help of VR requires a design in close analogy to natural conditions. With navigation in VR, most of the sensory input lies in the visual domain. Therefore, we aimed in our study at providing a large virtual environment with considerable detail of visual information with a fully immersive head-mounted setup to support immersion and a close-to-real-world experience. However, spatial updating also depends on vestibular and kinesthetic information (Chance, Gaunet, Beall, & Loomis, 1998; Klatzky, Loomis, Beall, Chance, & Golledge, 1998; Loomis et al., 1993; Riecke, Cunningham, & Bülthoff, 2007; Wang & Spelke, 2000). The performance of spatial tasks has been shown to improve with active exploration and natural movement in the environment (Klatzky et al., 1998; Mou & McNamara, 2002; Waller et al., 2004). As free-space walking requires considerably more space, improved tracking systems, and increased safety management, we realized movements in our setup with joystick translations and bodily rotations on a swivel chair in analogy to previous work (Riecke et al., 2010). Riecke et al. (2010) found with an adapted design of Ruddle and colleagues (Ruddle & Lessels, 2006; Ruddle & Lessels, 2009; Ruddle, Volkova, & Bülthoff, 2011) that full-body rotations already resulted in comparable performance levels to full walking. This is supported by real-world studies, which discovered a higher accuracy in direction estimation with physical body turns than imagined turns (Mou et al., 2004; Presson & Montello, 1994; Rieser, 1989). In contrast, Ruddle and colleagues (Ruddle & Lessels, 2006; Ruddle & Lessels, 2009; Ruddle, Volkova, & Bülthoff, 2011) found that translational movement was more important than body turns alone for improved accuracy in spatial tasks (e.g., cognitive maps measured by direction and straightline distance estimations or search tasks) with virtual environment exploration. Thus, for a fully immersive embodied experience in VR, free-space walking would be desirable to include all movement aspects as close to real-world sensory experience. Nevertheless, providing high visual detail and naturalistic visual flow and adding more naturalistic movement by bodily rotations giving kinesthetic and vestibular information to spatial navigation research in virtual environments reduces the gap between classical lab and real-world conditions and makes it possible to investigate embodied aspects of spatial cognition.

The importance of understanding results derived in virtual environments and the extent to which they can be related to natural conditions also derives from methodological limitations to the investigation of brain activity. Investigation of brain areas that are involved in spatial navigation (Bellmund, Gärdenfors, Moser, & Doeller, 2018; Chadwick, Jolly, Amos, Hassabis, & Spiers, 2015; Doeller, Barry, & Burgess, 2010; Ekstrom, 2010; Ekstrom, Huffman, & Starrett, 2017; Epstein, Patai, Julian, & Spiers, 2017; Gramann et al., 2010; Maguire, Nannery, & Spiers, 2006; Shine, Valdés-Herrera, Tempelmann, & Wolbers, 2019; Spiers & Maguire, 2007; Zhang, Copara, & Ekstrom, 2012) faces the problem that investigations of physiological processes in humans are mainly performed with fMRI, PET, MEG, or EEG. For the former three methods, human participants are required to lie or sit still and passively view a virtual environment. Thus, the results of brain activation during navigation often derive from VR experiments without physical movements. Recording brain activity during spatial navigation in a virtual environment with full mobility would be a desirable solution for future research. Specifically for straight-line walking, reliable and validated implementations are already available (Gwin, Gramann, Makeig, & Ferris, 2011; Oliveira, Schlink, Hairston, König, & Ferris, 2016). Recording in a natural environment allows the investigation of much more complex movement patterns and their influences on cognitive processes (Reiser, Wascher, & Arnau, 2019). Recent research including natural movement indicates that the lack of physical movement in simple virtual environments leads to quantitatively and qualitatively different physiological processes (Bohbot, Copara, Gotman, & Ekstrom, 2017; Ehinger et al., 2014). Ehinger et al. (2014) investigated brain activity with mobile EEG during a spatial path integration task to gain insight into brain activity during real-world navigation. They found significant differences in alpha activity in cortical clusters with respect to passive or active movement conditions and thus the availability of vestibular and kinesthetic input, respectively. Taken together, investigation in VR is of high importance for spatial navigation research and investigation of brain activity in humans. Further research is needed to determine the sweet spot for studies of spatial cognition of ecological validity and experimental control on the broad range of simple laboratory setups and VR setups of different degrees of sophistication and real-world conditions.

## Acknowledgments

We gratefully thank Jasmin Walter, Lara Syrek, Lucas Essmann, Paula Eisenhauer, Valerie Meyer, Carla S. Lembke, Timo Forbrich, Raul Sulaimanov and Marketa Becevova for technical assistance. Funded by the Deutsche Forschungsgemeinschaft (DFG, German Research Foundation) - project number GRK-2340/1 (DFG Research Training Group Computational Cognition) (AK, PK). Funded by the Deutsche Forschungsgemeinschaft (DFG, German Research Foundation) - project number GRK-2185/1 (DFG Research Training Group on Situated Cognition) (NK, PK). Funded by ErgoVR (BMBF Call: KMU innovative: Technologiebereich Mensch-Technik-Interaction) - joint research project number V5KMU17/221 (AK, PK). We gratefully acknowledge the support by the Open Access Publishing Fund of Osnabrück University.

